# Spinal motor neuron development and metabolism are transcriptionally regulated by Nuclear Factor IA

**DOI:** 10.1101/2024.06.26.600888

**Authors:** Julia Gauberg, Kevin B. Moreno, Karthik Jayaraman, Sara Abumeri, Sarah Jenkins, Alisa M. Salazar, Hiruy S Meharena, Stacey M Glasgow

## Abstract

Neural circuits governing all motor behaviors in vertebrates rely on the proper development of motor neurons and their precise targeting of limb muscles. Transcription factors are essential for motor neuron development, regulating their specification, migration, and axonal targeting. While transcriptional regulation of the early stages of motor neuron specification is well-established, much less is known about the role of transcription factors in the later stages of maturation and terminal arborization. Defining the molecular mechanisms of these later stages is critical for elucidating how motor circuits are constructed. Here, we demonstrate that the transcription factor Nuclear Factor-IA (NFIA) is required for motor neuron positioning, axonal branching, and neuromuscular junction formation. Moreover, we find that NFIA is required for proper mitochondrial function and ATP production, providing a new and important link between transcription factors and metabolism during motor neuron development. Together, these findings underscore the critical role of NFIA in instructing the assembly of spinal circuits for movement.

## INTRODUCTION

The spinal cord of vertebrates is composed of neuronal circuits that facilitate complex motor behaviors and dexterity. Human neurodevelopmental disorders that affect movement are either fatal or extremely debilitating, and most lack therapies or cures ^1^. Establishment of proper connectivity begins during development, when motor neurons migrate within the spinal cord and innervate peripheral targets. Transcription factors play a critical role in regulating motor neuron identity, positioning within the spinal cord, and connectivity patterns along the body axis ^1–8^.

Transcription factors essential for early motor neuron specification from progenitors (e.g LIM, Hox, Fox, Homeodomain, Onecut) are well-characterized ^9,10^. In contrast, little is known about the transcriptional codes that regulate post-mitotic motor neuron maturation, which involves acquiring terminal identity features, defining innervation patterns, and forming synapses.

The formation of connections between motor neurons and their target muscles follows acquisition of motor neuron identity and migration within the spinal cord. Motor neurons undergo several iterations of clustering such that groups of motor neurons that target related areas are clustered together into columns and pools within the ventral spinal cord ^9^. In thoracic regions of the spinal cord, median motor columns (MMC), which target axial muscles, form along with preganglionic (PGC) neurons that target the diaphragm, and hypaxial motor columns (HMC) that target abdominal muscles. Motor neurons at limb-level regions of the spinal cord are hierarchically segregated into MMC and lateral motor columns (LMC). The LMC column is further divided into medial (LMCM) and lateral sections (LMCL), which innervate the ventral and dorsal limb, respectively ^11–13^. Next, LMC motor columns undergo a secondary reorganization into sub-clusters consisting of motor pools, which are compact groups of motor neurons that share intrinsic characteristics and establish distinct axon projection patterns ^14–19^. Although extensive research has been conducted on the medio-lateral and dorso-ventral organization of motor neurons, the precise molecular and cellular programs coordinating the ordered migration of motor neurons to the ventral horn and their sorting into motor pools are not completely understood.

Once motor neurons are organized in the spinal cord, they begin to send motor axons out towards their targets, such as the limb. Within the limb mesenchyme, these axons further organize into main nerve trunks, each following distinct pathways that are dependent on the identity of the motor neuron^20^. Along these trunks, smaller clusters of motor axons branch off at specific locations, forming finer nerve branches that eventually reach their designated target muscles in a process called terminal arborization ^21–23^. In some disease models, such as Spinal Muscular Atrophy, initiation of motor axon outgrowth and pathfinding is normal, but axonal branching and neuromuscular junction formation is disrupted ^24,25^. This suggests that the processes of axonal branching and terminal arborization are distinct from initial axonal outgrowth and rely on different mechanisms ^26^. While the relationship between motor neuron identity and initial axon trajectories is well established, the molecular mechanisms that co-regulate motor neuron identity, axonal branching, and terminal arborization, which occur later in development, are not completely understood.

Not only is axonal branching tightly controlled mechanistically, it is also an energetically expensive process. Therefore, axon extension necessitates the proper function of mitochondria, which supply energy to growing and branching axons ^27,28^. When energy demands are not met, axons fail to extend and branch ^29,30^. However, the molecular mechanisms that regulate energy supply during development remain poorly understood. Furthermore, how transcription factors regulate energy metabolism in developing motor neurons is not known.

Our previous investigation into Nuclear Factor IA (NFIA) function in the spinal cord revealed that NFIA may be a key player in post-mitotic motor neuron development ^31^. This is consistent with a recent omic study, which revealed that NFIA is expressed in a temporally-dynamic fashion in motor neurons ^32^. However, the function of NFIA in this neuronal population is unknown.

Here, we identified the transcription factor NFIA as an essential regulator of the development and differentiation of limb-level (brachial and lumbar) motor neurons. Using a multi-omic approach we defined cellular and molecular processes directly regulated by NFIA in the developing spinal cord. These analyses identified several biological processes associated with cellular metabolism, cell positioning, and axon projections that are regulated by NFIA activity. Functional analyses revealed that NFIA regulates limb-innervating motor neuron positioning and identity in the ventral spinal cord. In the absence of NFIA initial axonal trajectories were normal, but branching and formation for neuromuscular junctions (NMJs) was compromised, indicating that NFIA plays a critical role in these later processes. Finally, we observed that NFIA regulates motor neuron cellular metabolism, revealing previously unappreciated functions of NFIA in the CNS. This work provides fundamental insights into mechanisms of developmental neuronal diversification, cell organization, and motor neuron circuit assembly.

## RESULTS

### NFIA is expressed in spinal motor neurons throughout development

In the developing mouse spinal cord, postmitotic motor neurons are generated between E9 and E11 ^4^. To characterize NFIA expression during spinal motor neuron development we first examined mouse spinal cords at limb and thoracic levels between day E10.5 and postnatal day P4. NFIA is first detectable in the mouse spinal cord at e11.5 (Figure 1A) after motor neurons have been specified from progenitors. At this time, NFIA is expressed in (1) the motor neuron progenitor (pMN) domain, (2) motor neurons emerging from the pMN domain, and (3) laterally positioned post-mitotic motor neurons (Figure 1A, arrows). NFIA expression in motor neurons persists throughout development and is maintained postnatally (Figure 1A). To determine which motor columns express NFIA we compared its expression with transcription factor markers of limb-level columns (MMC and LMC) and thoracic-level columns (MMC, HMG and PGC) at E13.5. In brachial-level motor neurons, NFIA is co-expressed with Isl1/2,FoxP1 (Figures 1 B-D and S1A-D), and Hb9 (Figure 1E-G) transcription factors. FoxP1 and Isl1/2 are expressed in both LMCL and LMCM motor neurons, whereas Hb9 demarcates LMCL motor neurons. In MMCs, NFIA is co-expressed with Hb9 (Figure 1E-G) and Lhx3 (Figure 1H-J). At thoracic levels NFIA is expressed in HMC and MMC motor columns, distinguished by Isl1/2 (Figure 1K-M) and Hb9 (Figure 1N-P) co-expression. In contrast, NFIA is excluded from thoracic PGC neurons (Figure 1N-P gray circle). Taken together, NFIA is found in MMC, LMCL, LMCM, and HMC motor neurons, but is absent from thoracic PGC neurons (Figure 1Q).

**Figure 1.**
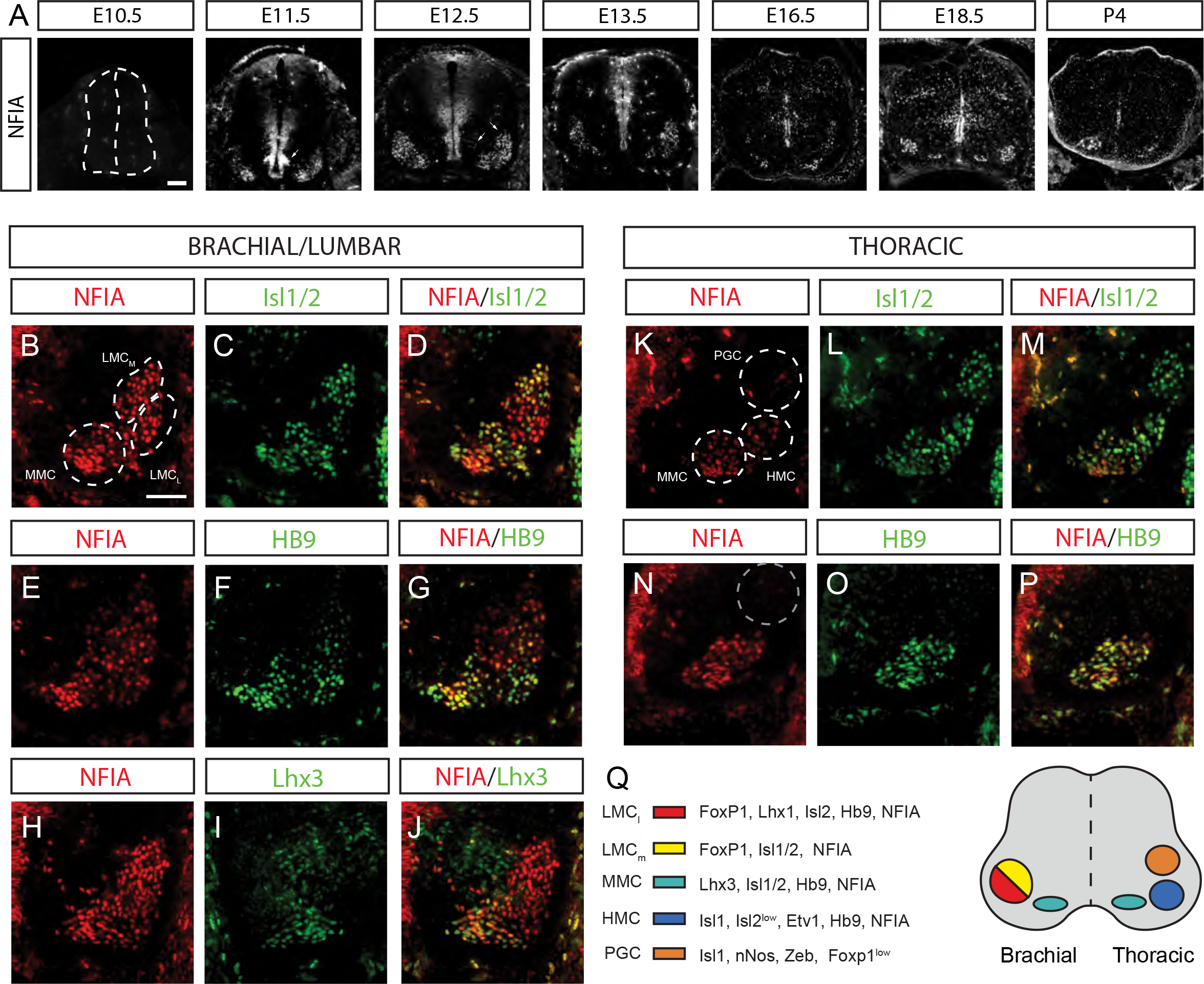
NFIA is expressed in spinal motor neurons from E11.5 to P4 (A) Patterns of NFIA expression in wildtype limb-level spinal cord from E10.5 to P4. Arrows indicate the motor neuron progenitor domain (E11.5), as well as motor neurons emerging from the pMN (e12.5), and laterally positioned post-mitotic motor neurons (E12.5) (B-J) Immunofluorescent staining of NFIA with motor column markers within brachial/lumbar spinal cord regions at E13.5. Right ventral quadrant of the spinal cord is shown. Dashed circles indicate the median motor column (MMC) and two lateral motor columns (LMC): medial (LMCM) and lateral (LMCL) (K-P) Immunofluorescent staining of NFIA with motor column markers within the thoracic spinal cord at E13.5. Right ventral quadrant of the spinal cord is shown. Dashed circles indicate the MMC, hypaxial motor column (HMC) and preganglionic motor column (PGC). Dashed gray circle in panel (N) highlights the absence of NFIA in the PGC (Q) Summary of NFIA expression with other known motor column markers in E13.5 mouse spinal cord. Scale bars represent 100 μm.

### NFIA directly regulates genes required for spinal motor neuron development

While the function of NFIA as a glial determining factor in the CNS has been well documented ^33–35^, its function in neuronal populations is less clear, and the mechanisms of NFIA activity in motor neurons are undefined. To understand the mechanism of NFIA action in developing spinal neurons, we generated CNS-specific NFIA conditional knockout mice, in which NFIA was deleted by Nestin-Cre (NFIA^f/f^;Nestin-Cre, referred to as NFIAcKO). NFIA protein expression was largely eliminated from developing spinal motor neurons as marked by co-expression with Isl1 at brachial- and thoracic- levels (Figure S2A-F). NFIA remained expressed in a small population of motor neurons, possibly signifying that these cells started expressing NFIA prior to Cre expression. Given that another Nuclear factor-I family member, NFIB, has been shown to be expressed in developing motor neurons ^32,35–38^, we first evaluated whether loss of NFIA results in changes in NFIB expression. We found that the number of NFIB-positive cells was unchanged in NFIAcKO motor neurons at brachial, lumbar, and thoracic levels (Figure S2G-L), indicating that NFIB levels do not compensate for NFIA loss in these cells.

To examine the mechanistic role of NFIA in developing neurons of the spinal cord, we performed RNA-seq on control (NFIA^f/f^) and NFIAcKO spinal cords at E13.5. This identified 2,511 differentially expressed genes (DEGs) in the absence of NFIA, with a threshold fold change |FC|>0.3 and adjusted P<0.05 (Figure 2A and S3A; Table S1). Of these, 55% were upregulated and 45% were downregulated. To validate these results, we performed quantitative polymerase chain reaction (qPCR) for NFIA, revealing a significant reduction of NFIA in NFIAcKO spinal cords (Figure S3B). In addition, we validated a subset of highly-expressed genes that were downregulated in NFIAcKO spinal cords, recapitulating the expression patterns observed via RNA-seq analysis (Figure S3B).

**Figure 2.**
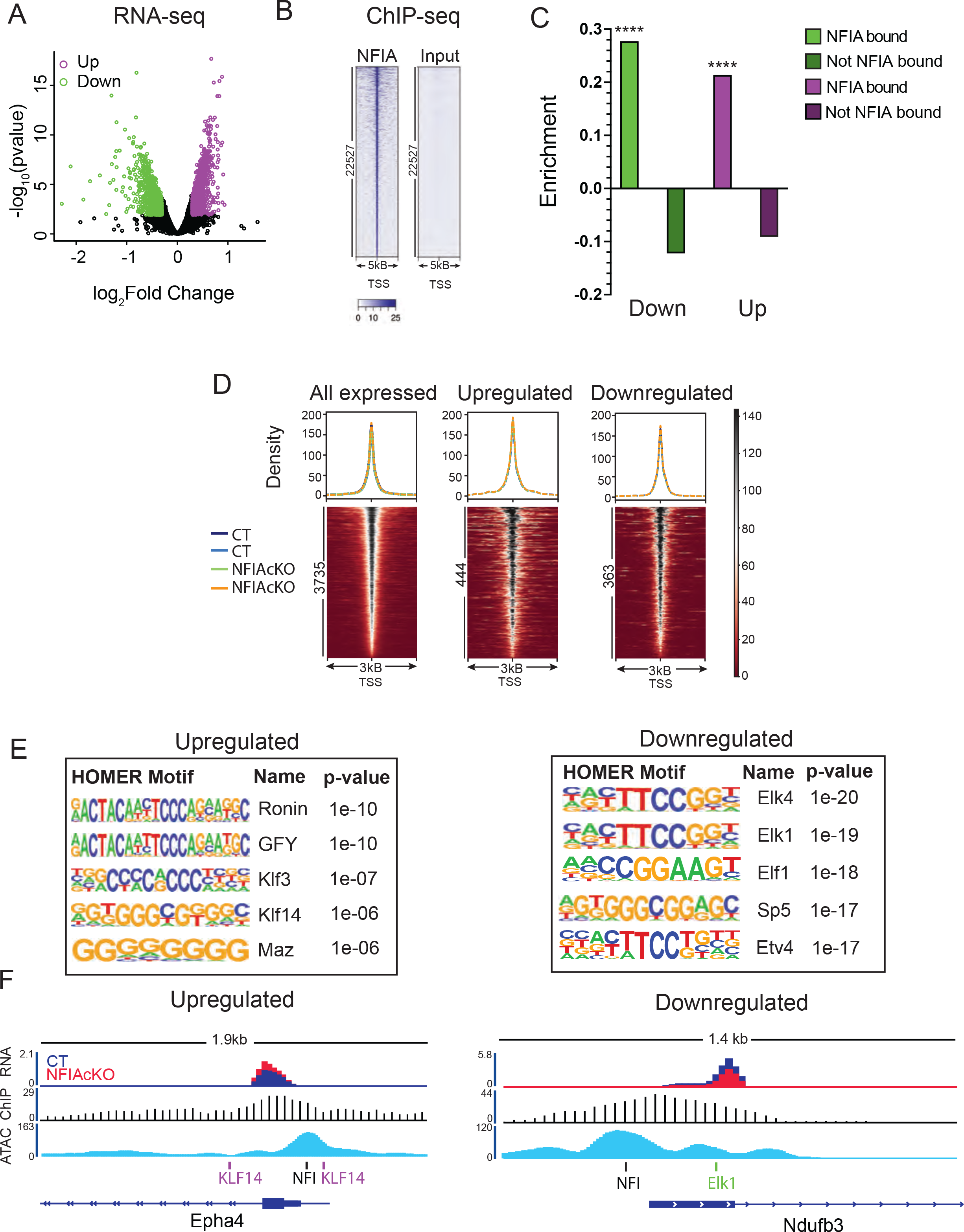
NFIA directly regulates expression of genes in E13.5 spinal cords (A) Volcano plot depicting differential gene expression between control (CT) and NFIA conditional knockout (NFIAcKO) spinal cords (n=3) (B) NFIA ChIP-seq signals at NFIA-bound genomic regions in CT spinal cords (C) Enrichment plot of down- and upregulated genes that are bound versus not bound by NFIA at their promoters. p**** ≤0.0001 as determined by a randomized permutation test (D) Heatmap coverage plot of accessible chromatin in promoters of CT and NFIAcKO motor neuron genes. All expressed, upregulated, and downregulated genes that were bound by NFIA were examined (n=2) (E) Top enriched motifs at NFIA-bound promoters of upregulated and downregulated genes in CT motor neurons (F) RNA-seq, ATAC-seq, and ChIP-seq gene tracks around the promoter of one upregulated (left) and one downregulated (right) gene. Vertical bars depict NFI, Klf14, and Elk1 transcription factor motifs in this region.

To define the DEGs that are directly regulated by NFIA we performed chromatin immunoprecipitation followed by sequencing (ChIP-seq) from E13.5 spinal cords. This profiling identified 19,655 NFIA binding sites, which were enriched near the TSS (± 5 kb) of genes (Figure 2B; Table S2). The majority of peaks mapped to intronic regions (36%), followed by intergenic (32%) and promoter (14%) regions (Figure S3C). These data suggest that NFIA not only controls transcription through proximal promoters, but likely regulates a larger portion of genes via distal enhancers. We extracted the ChIP-seq peaks within promoter regions (2kb upstream and 500bp downstream) and identified 4,453 expressed genes that were bound by NFIA on their promoters (Figure S3D). Of these, 447 were upregulated and 376 were downregulated in NFIAcKO spinal cords (Figure S3D). Next, we performed a randomized permutation test (10^4^) and observed a significant enrichment of both up- (pValue <10^-4^) and down-regulated genes (pValue <10^-4^) with NFIA occupancy on the promoter. On the other hand, genes not bound by NFIA had no statistically significant enrichment for DEGs (Figure 2C), indicating that NFIA is the primary driver of the altered transcriptional state.

We hypothesized that the loss of NFIA induces dysregulation of gene expression through the binding of other transcription factors to accessible promoters of motor neuron DEGs. To explore this, we first identified accessible chromatin regions in E13.5 motor neurons. We sorted control and NFIAcKO motor neurons expressing the Hb9::GFP transgene using fluorescence activated cell sorting (FACS; Figure S3E) and performed Assay for Transposase-Accessible Chromatin using sequencing (ATAC-seq; Table S3) . We then compared accessibility in control and NFIAcKO motor neurons and, surprisingly, discovered that there was no difference in ATAC-seq peaks between control and NFIAcKO motor neurons (Figure 2D).

Next, we isolated ATAC-seq peaks within the promoters of genes harboring NFIA (3,735 peaks), to explore other transcription factor binding sites associated with DEGs. To determine which transcription factors may be co-regulating expression of NFIA target genes, we performed motif enrichment analysis using HOMER ^39^ on accessible promoters harboring NFIA. We observed that up- and down-regulated genes showed distinct motif enrichment patterns in their promoters (Figure 2E). Promoters of upregulated genes are enriched for transcription factor motifs, such as the Thanatos Associated Proteins (THAP) and Sp, and Krüppel-like factor (KLF) families of transcription factors. In contrast, NFIA-harboring downregulated genes harbor motifs of the ETS family of transcription factors, such as ELK1. Motifs bound by these factors could be seen in the vicinity of NFIA ChIP-seq peaks within promoters (Figure 2F). Therefore, in the absence of NFIA, these transcription factors may bind to the same open chromatin regions and modulate expression of NFIA-target genes. These results suggest that NFIA is the dominant driver of gene expression necessary for proper maturation of motor neurons.

To remove inaccessible genes from our DEG list, we integrated RNA-seq, ChIP-seq, and ATAC-seq data, revealing 355 downregulated and 422 upregulated genes (Figure 3A-B and Table S4). GO term analysis revealed that upregulated genes were significantly enriched for neuron development and projection morphogenesis, whereas downregulated genes were enriched for processes related to translation and mitochondrial respiration (Figure 3C).

**Figure 3.**
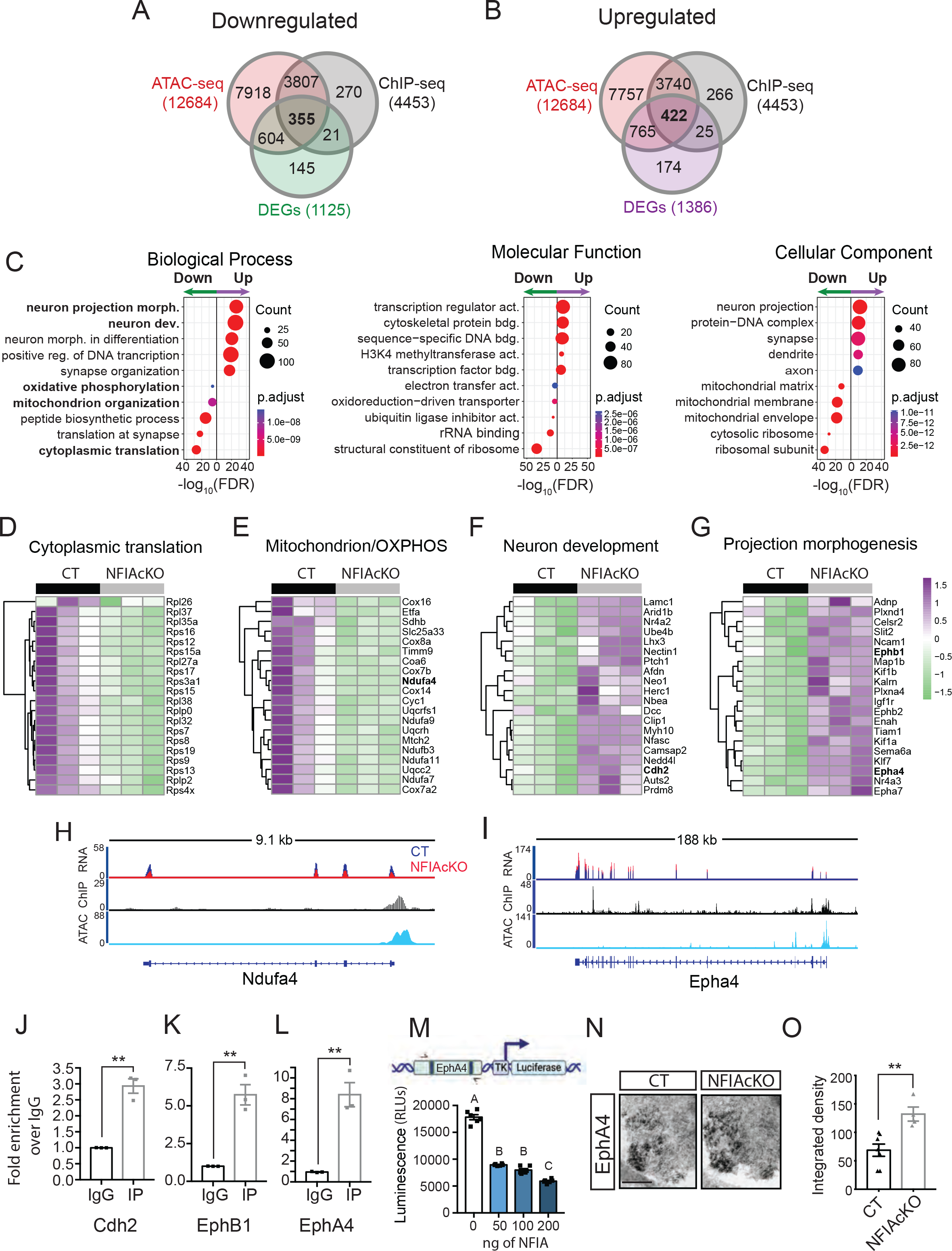
NFIA directly regulates genes required for spinal motor neuron development (A-B) Venn diagrams showing overlap between NFIA-bound promoters, accessible promoters in motor neurons, and genes either downregulated (A) or upregulated (B) in NFIA conditional knockouts (NFIAcKO) (C) Gene ontology (GO) analysis of NFIA-bound DEGs in control (CT) versus NFIAcKO motor neurons falling under the categories of biological process, molecular function, and cellular component. GO analysis was performed separately on up- and downregulated genes (D-G) Heatmap and hierarchical clustering of top 20 genes from bolded categories in (C), showing differences between CT and NFIAcKO gene expression. Bolded gene names are further examined in (H-O) (H-I) RNA-seq, ATAC-seq, and ChIP-seq gene tracks around the Ndufa4 (H) and Epha4 (I) loci. (J-L) ChIP-qPCR analysis for Cdh2 (A), Ephb1 (B), and Epha4 (C) using antibodies against NFIA or mouse control IgG in CT spinal cords (n=3) (M) Luciferase reporter assay of the Epha4 promoter in the presence of 50, 100, and 200 ng of NFIA in HEK293 cells. RLU= relative light unit (n=6) (N) Representative in-situ hybridization image of Epha4 within motor neurons of CT and NFIAcKO spinal cords (O) Quantitative comparison of Epha4 in-situ hybridization signal between CT and NFIAcKO motor neurons (n=4-7). All data are represented as mean ± SEM. p* ≤0.05; p** ≤0.01 by Student’s T-test. Different uppercase letters signify p≤0.001 as determined by a One-Way ANOVA followed by Tukey’s test.

Heatmaps of the top 20 genes in each category identified multiple downregulated ribosomal (e.g. Rpl26, Rpl37, Rps8, Rps17) and oxidative phosphorylation genes (e.g. Ndufa4, Ndufb3, Sdhb; Figure 3D-E), as well as upregulated genes related to neuronal migration and adhesion (e.g. Cdh2, Afdn, Nectin1) and axon guidance (e.g. Epha4, Ephb1, Plxna4; Figure 3F-G).

Visualization of select RNA-, ChIP- and ATAC-seq peak diagrams revealed that accessible promoter regions of axonal and metabolism related genes, such as Ndufa4, Epha4, Ephb1 and Cdh2, are bound and regulated by NFIA (Figure 3H-I and S3F). NFIA occupancy was validated on E13.5 mouse spinal cords with chromatin-immunoprecipitation followed by qPCR (ChIP-qPCR) for a subset of genes including: Cdh2, Ephb1, and Epha4 (Figure 3J-L). Using EphA4 as an example, we analyzed the ChIP-seq defined NFIA occupancy region nearest the promoter, finding two predicted NFIA binding sites. To confirm NFIA regulation of EphA4, we performed a luciferase reporter assay with the EphA4 promoter region and found that NFIA could transcriptionally repress luciferase activity (Figure 3M). To determine if this regulation was functionally relevant, we examined Epha4 mRNA levels in E13.5 control and NFIAcKO motor neurons, observing that Epha4 mRNA levels were elevated in the absence of NFIA (Figure 3N- O). Overall, these results demonstrate that NFIA directly regulates the expression of genes related to translation, oxidative phosphorylation, neuron migration, and neuron projection morphogenesis in the spinal cord of developing mice.

### NFIA mutants exhibit impaired divisional segregation of limb-level motor columns and pools

Given that our multi-omic data revealed that NFIA directly regulates genes related to motor neuron development, migration, and morphology (e.g. Cdh2, Nectin1, Epha4, Ephb1), we sought to determine the functional consequences of NFIA deletion on motor neuron development. Although NFIA expression is initiated in motor neurons after the onset of specification (Figure 1A), we tested the impact of NFIA activity on motor neuron progenitors. Consistent with the timing of NFIA expression, the number of Olig2+ progenitor cells and Nkx2.2+ cells, the latter located in the p3 domain just ventral to the pMN, were not affected in NFIAcKO mice (Figure S4). Next, we asked whether loss of NFIA leads to gross changes to developing motor neurons. At E13.5 the number of Hb9^+^, Isl1^+^, Lhx1^+^, and Foxp1^+^ cells were not significantly altered between NFIAcKOs and control motor neurons (Figure 4 and S5A-D).

**Figure 4.**
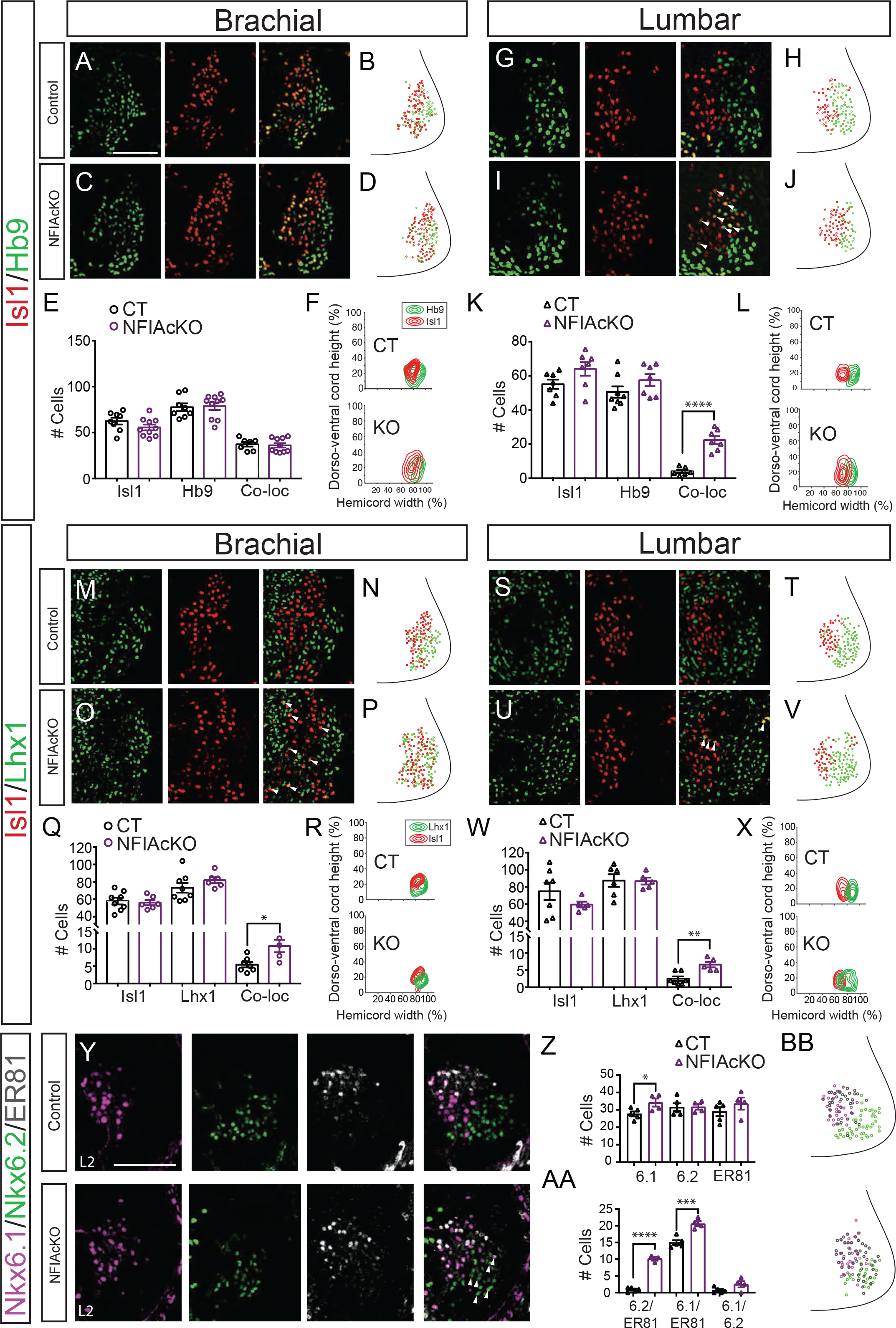
NFIA mutants exhibit impaired divisional segregation of limb-level motor columns and pools (A and C) Immunofluorescent staining of LMC markers Isl1/Hb9 on control (CT; NFIA^f/f^) and NFIA conditional knockout (NFIAcKO; NFIA^f/f^;Nestin-cre) E13.5 ventral spinal cords at brachial levels (B and D) Digitally reconstructed distribution of CT (B) and NFIAcKO (D) LMC neurons at brachial level, shown as a transverse projection (dot plot) (E) Quantification of Isl1-positive, Hb9-positive, and Isl1/Hb9 co-positive (Co-loc) cells in LMC neurons in CT and NFIAcKO spinal cords at brachial level. (n=7-10) (F) Transverse contour density plots of LMCM (Red-Isl1) and LMCL (Green-Hb9) neurons in CT and NFIAcKO ventral spinal cord (G-L) Distribution analysis of LMCM (Red-Isl1) and LMCL (Green-Hb9) in CT and NFIAcKO E13.5 ventral spinal cords at lumbar level. Arrowheads indicate Isl1/Hb9 co-positive cells in NFIAcKO spinal cords. (n=6-8) |(M and O) Immunofluorescent staining of LMC markers Isl1/Lhx1 on CT and NFIAcKO ventral spinal cords at brachial levels (N and P) Digitally reconstructed distribution of Ct (N) and NFIAcKO (P) LMC neurons at brachial level, shown as a transverse projection (dot plot) (Q) Quantification of Isl1-positive, Lhx1-positive, and Isl1/Lhx1 Co-loc cells in LMC neurons in CT and NFIAcKO spinal cords at brachial level. (n=6-8) (R) Transverse contour density plots of LMCM (Red-Isl1) and LMCL (Green-Lhx1) neurons in CT and NFIAcKO ventral spinal cords (S-X) Distribution analysis of LMCM (Red-Isl1) and LMCL (Green-Lhx1) in CT and NFIAcKO ventral spinal cords at lumbar level. Arrowheads indicate Isl1/Lhx1 co-positive cells in NFIAcKO spinal cords. (n=5-7) (Y) Immunofluorescent staining of motor pool markers Nxk6.1/Nkx6.2/Er81 at L2 lumbar level in E13.5 CT (top) and NFIAcKO (bottom) spinal cords (Z-AA) Quantification of L2 neurons in CT and NFIAcKO lumbar spinal cord. (n=3-5) (BB) Digitally reconstructed distribution of CT (top) and NFIAcKO (bottom) lumbar L2 motor pools, shown as a transverse projection (dot plot). For all images, right ventral quadrant of the spinal cord is shown. Scale bars represent 100 μm. All data are represented as mean ± SEM. p* ≤0.05; p** ≤0.01, p***≤0.001 , p****≤0.0001 by Student’s T-test.

Furthermore, no changes were observed in cell death as marked by activated Caspase3 (Figure S5E-J), indicating that the total number of motor neurons is unchanged in NFIAcKO mice.

Together, these data suggest that loss of NFIA does not affect progenitor populations, patterning of the ventral spinal cord ventricular zone, or the overall number of motor neurons.

To further investigate how NFIA is contributing to motor neuron development, we analyzed the expression of markers for motor neuron identity and columnar differentiation in NFIAcKO mice, focusing on brachial and lumbar spinal cord levels in E13.5 embryos. In the MMC, markers Hb9 and Isl1 were unaffected, indicating that this population’s identity may not be reliant on NFIA expression (Figure S5K-L). Next, we examined more laterally located LMCs with two combinations of markers Isl1/Hb9 and Isl1/Lhx1, which demarcate the subdivision between LMCM (Isl1) and LMCL (Hb9 and Lhx1). At brachial and lumbar levels, no change in Isl1^+^, Hb9^+^, or Lhx1^+^ cell number was observed (Figure 5A-W), however, the spatial distribution of LMC cell bodies was altered. LMC neurons normally complete inside-out radial migration by E13.5 ^13,40,41^. However, LMCM (Isl1^+^) cells were abnormally distributed and shifted laterally in NFIAcKO, invading the LMCL space and intermingling with Hb9^+^ cells (Figure 4D-F, J-L-).

**Figure 5.**
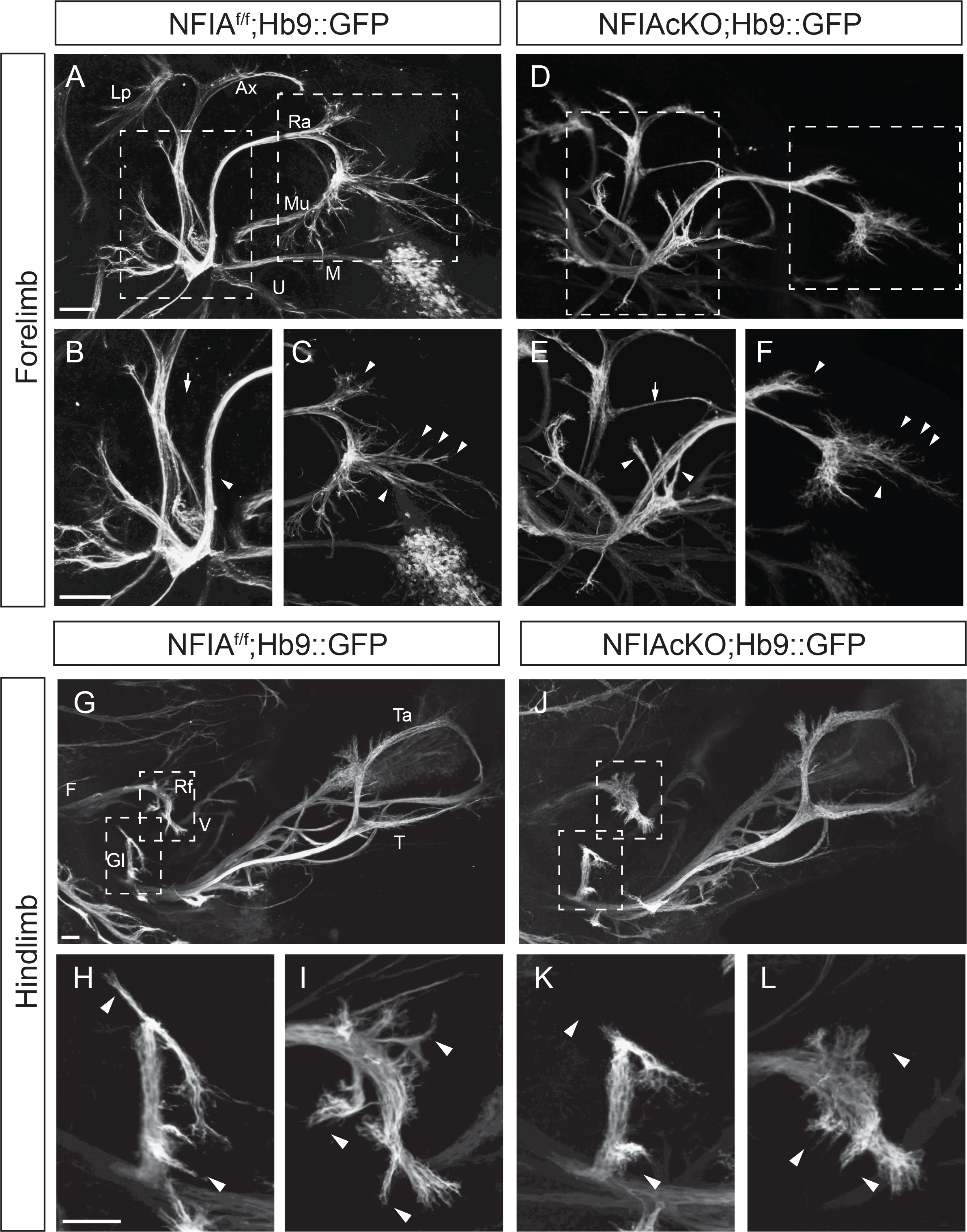
Aberrant limb innervation in the absence of NFIA (A and D) Optically-cleared whole mount GFP staining of forelimb innervation patterns in control (A; NFIAff;Hb9::GFP) and NFIAcKO (B; NFIAff;Nestin-Cre;Hb9::GFP) embryos at E13.5 (n=4- 5) (B-C and E-F) Higher magnification images of radial and axillary nerves in control (B-C) and NFIAcKO (E-F) forelimbs. Arrowheads indicate major nerve defects. (G and J) Whole mount GFP staining of hindlimb innervation patterns in control and NFIAcKO embryos at E13.5. (H-L) Higher magnification images of gluteus and vasti muscle nerves in control (H-I) and NFIAcKO (K-L) hindlimbs Abbreviations: Axillary (Ax), Gluteus (Gl), Lateral pectoral (Lp), Median (M), Musculocutaneous (Mu), Radial (Ra), Rectus femoris (Rf), Tibialis (T), Tibialis anterior (Ta) Ulnar (U), Vasti (V) Arrows and arrowheads indicate major nerve defects. Scale bars represent 100μm.

Lhx1^+^-LMCL cells were similarly intermingled with Isl1^+^-LMCM cells in both brachial and lumbar spinal cord (Figure 4M-X). Interestingly, a small subset of cells expressed markers of both LMCL and LMCM populations, suggesting that these cells had a mixed identity or failed to properly segregate (Figure 4A-X arrowheads). Taken together, NFIA is not required for the initial specification of motor neurons, but is involved in the distribution of LMC cells and maintaining LMC boundary integrity.

Since limb-innervating motor neurons are further organized into motor pools, we examined whether NFIA is required for pool clustering and segregation. We focused on motor pools at the L2-L3 level of the lumbar spinal cord because they have defined populations with dorso-ventral and medial-lateral divisions ^6,19^. In this region, NFIA is expressed in L2/L3 motor pools and is markedly reduced in NFIAcKO spinal cords (Figure S6A-B). At L2 Nkx6.1+/Er81+ define the adductor/gracilis (A/G)-innervating pool, Nkx6.2 marks the femoris/tensor fasciae latae (R/T)-innervating pool, Nkx6.1 defines the adductor brevis (Ab)-innervating pool, and the vasti (V)-innervating pool is labeled by Er81 expression (Figure S6C). In the absence of NFIA we found a modest overall increase in the number of Nkx6.1+ cells at the L2 level (Figure 4Y-Z) and an increase in Nkx6.1+ and Er81+ cells at the L3 level (Figure S6D-F). A/G-innervating pools, co-labeled by Nkx6.1+/Er81+, were increased in NFIAcKO spinal cords, consistent with the rise in both Nk6.1+ and Er81+ cells (Figure 4AA and S6G). In control spinal cords, R/T- and V-innervating pools are clustered in discrete locations with nominal co-expression of Nkx6.2 and Er81 (Figure 4Y and S6D). However, in NFIAcKO embryos, Er81 expression was more dispersed and displayed significant co-expression with Nkx6.2 (Figure 4AA-BB and S6F-I), indicating that there was intermingling of these LMC divisions. Overall, NFIA controls segmentation of limb-innervating motor neurons at both the columnar and motor pool levels, which is consistent with the molecular changes at the RNA level observed using RNA-seq.

### Aberrant axonal projections, arborization, and neuromuscular junctions in the absence of NFIA

Since NFIA-bound DEGs were enriched for genes associated with neuronal projections (Figure 3), we reasoned that motor neuron pathfinding and/or terminal arborization may be compromised in the absence of NFIA. This line of reasoning is further supported by the fact that NFIAcKO embryos have defects in LMC motor neuron positioning and boundary formation (Figure 4), which has been shown to be critical for axonal pathfinding ^9,42^. Therefore, we analyzed the pattern of limb innervation by tracing axonal trajectories in E13.5 control and NFIAcKO embryos carrying the HB9::GFP transgene, which allows for the visualization of LMCL spinal motor axons. The pattern of eGFP-labeled nerve branches was highly reproducible in control embryos. Furthermore, in the absence of NFIA, bifurcation of LMC axonal projections to dorsal and ventral limb trajectories were largely normal (Figure S7). However, NFIA mutant embryos displayed aberrant axonal projections and marked defects in branching at distal regions of forelimbs and hindlimbs (Figure 5). Specifically, terminal branching patterns of the radial nerve were much less complex and had a stunted appearance, with decreased branch length in NFIAcKO embryos (Figure 5A, C, D, F; arrowheads). Additionally, NFIAcKO embryos also had aberrant radial nerve branches that were absent in control mice (Figure 5B,E; arrowheads) and the axillary nerve was connected to the radial nerve rather than the musculocutaneous nerve (Figure 5B,E; arrow). Similar innervation defects were found in hindlimbs of NFIAcKO embryos. Nerve branches targeting the gluteus and vasti muscles were irregular in the absence of NFIA (Figure 5G-L). Gluteus-targeting nerve branches were stunted or absent in certain trajectories (Figure 5H, K) and vasti-targeting nerve branches were reduced and disorganized (Figure 5I, L). Therefore, motor neurons exhibit a pronounced deficit in terminal axonal branching in NFIAcKO embryos.

To assess whether the axonal branching defects observed *in vivo* could also be detected outside the context of the limb (*ex vivo*), we cultured motor neuron explants from Hb9::GFP positive control and NFIAcKO E13.5 ventral spinal cords. Cells and out-growing axons expressed GFP in both control and NFIAcKO groups, indicating that they corresponded to spinal motor neurons, and anti-TUJ1 signal co-localized extensively with GFP (Figure S8A).

Axon outgrowth was reduced in both brachial (Figure 6A-C) and lumbar (Figure 6D-F) NFIAcKO motor neuron explants compared to controls.

**Figure 6.**
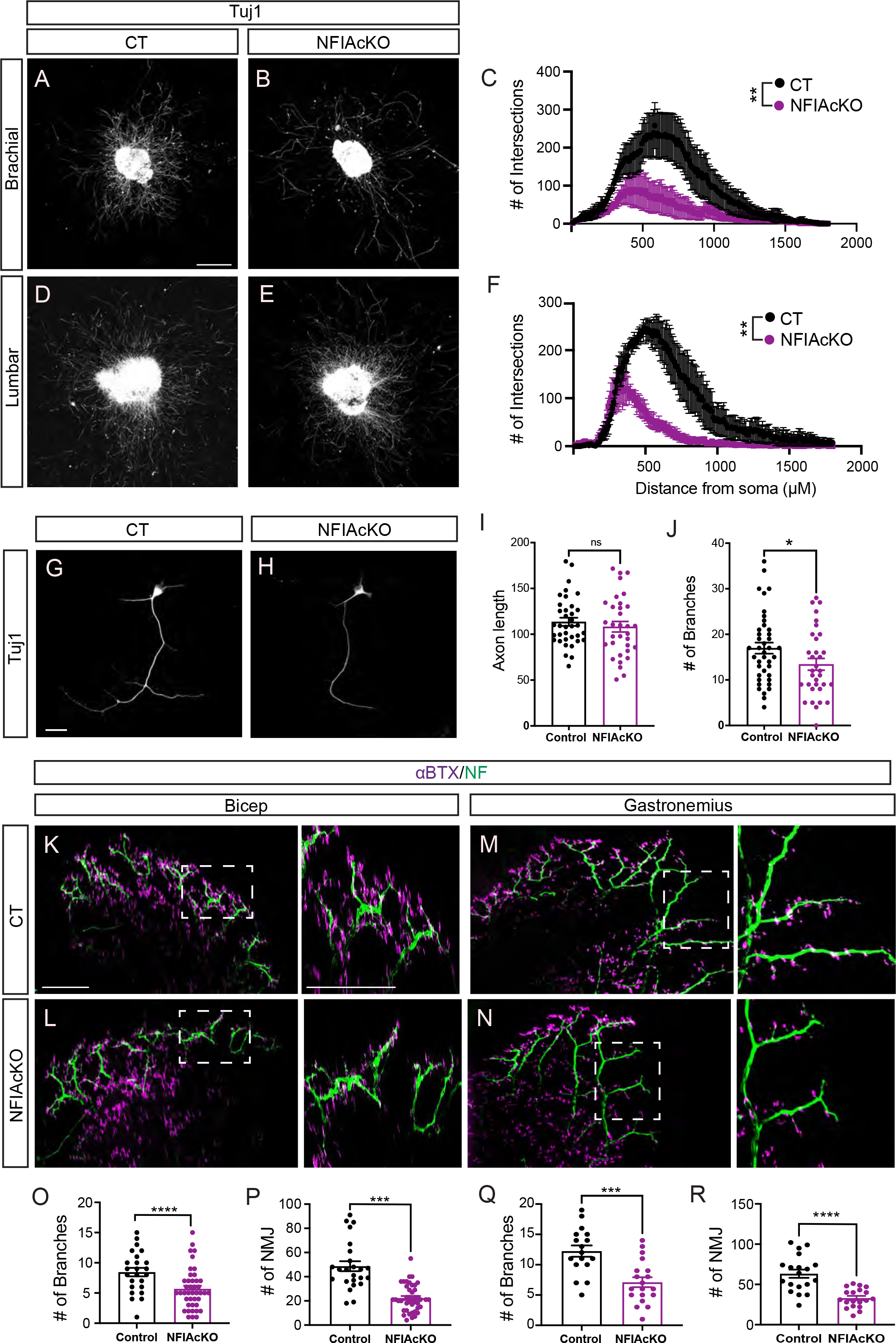
Defective axonal branching and neuromuscular junction formation in NFIA mutant mice (A,B and D, E) Representative images of neurite outgrowth in control (CT) and NFIA conditional knockout (NFIAcKO) explant cultures from E13.5 brachial and lumbar ventral spinal cords after 3-4 days in vitro (DIV). Scale bars represent 500 μm. (C and F) Scholl analysis of 3-4 DIV motor neuron explants from CT and NFIAcKO in brachial (C) and lumbar (F) regions. Analysis was performed with the ImageJ Scholl analysis plugin. Number of explant cultures analyzed were n=9 for brachial and n=6-7 for lumbar. (G-H) Representative images of motor neuron primary culture from E13.5 CT and NFIAcKO spinal cords. Scale bars represent 20μm. (I-J) Quantification of axon length (I) and number of branches (J) in primary motor neuron cultures. Data was obtained from a total of 33-40 motor neurons from 3-4 embryos. (K-L) Visualization of motor neurons and neuromuscular junctions (NMJs) in CT (K) and NFIAcKO (L) P0 bicep muscles using neurofilament (NF; green) and α-bungarotoxin (BTX; magenta). Boxed regions are shown at higher magnification on the right. Scale bars represent 200 μm. (M-N) Visualization of NMJs in CT (M) and NFIAcKO (N) gastrocnemius muscles. (O-P) Quantification of secondary axon branches and NMJ number in bicep muscles of CT and NFIAcKO mice. Data was obtained from a total of 17-19 motor neurons from 3-4 embryos. (Q-R) Quantification of secondary axon branches and NMJ number in gastrocnemius muscles of CT and NFIAcKO mice. Data was obtained from a total of 24-43 motor neurons from 3-4 embryos. All data are represented as mean ± SEM. p* ≤0.05; p** ≤0.01, p***≤0.001 , p****≤0.0001 by Student’s T-test.

Since NFIA is deleted from both neurons and glial cells in NFIAcKO embryos and explants, we cultured dissociated motor neurons *in vitro* to specifically determine if there are defects in axonal branching within this cell type. Control motor neurons expressed NFIA (Figure S8B) and minimal numbers of GFAP astrocytes were observed in primary motor neuron cultures (Figure S8C). Motor neuron axons from control and NFIAcKO ventral spinal cords were comparable in length (Figure 6G-I); however the NFIAcKO motor neurons were less complex, with fewer branches (Figure 6G-J), consistent with *in vivo* results.

Next, we wanted to determine if the disorganization of LMCs and the attenuation of axonal branching at E13.5 affected neuromuscular junction (NMJ) formation in the developing limbs. Since NFIA-mutant mice die at birth, we examined limb muscles at E18.5, a time when NMJs formation is actively occurring ^43,44^. Flat-mount preparations of NFIAcKO bicep (forelimb) and gastrocnemius (hindlimb) were co-stained for neurofilament (NF) and nicotinic acetylcholine receptors (using alpha-bungarotoxin; α-BTX). The axon bundles labeled by NF seemed largely normal in both bicep and gastrocnemius muscles of NFIAcKO mice, but secondary branches and terminal arbors were significantly altered in both muscle groups in the absence of NFIA. Specifically, the complexity of the secondary branches was reduced in NFIAcKO bicep muscles (Figure 6K-L,O) and the number of α-BTX nerve terminals were also reduced (Figure 6K-L,P).

Similar defects in hindlimb gastrocnemius muscles were observed (Figure 6M-N, Q-R). Altogether, NFIA mutant embryos display a reduction in axonal branching with compromised terminal arborization of skeletal muscle NMJs, suggesting that motor axons fail to fully extend within the limb in the absence of NFIA. Overall, early disruptions in muscle nerve branch formation are accompanied by later defects in axonal arborization and muscle innervation.

### Loss of NFIA leads to mitochondrial dysfunction in motor neurons

Differential gene expression analysis of accessible and NFIA-bound genes revealed a downregulation of genes involved in oxidative phosphorylation (e.g. Ndufs7, Nduf4a, Sdhb, Cyc1, Cox7B) in NFIAcKO spinal cords (Figure 3H and Figure 7A bold). In high-energy demanding neurons, mitochondria are essential for locally synthesizing ATP and maintaining energy homeostasis ^45^. It has also been well documented that mitochondrial dynamics play a critical role in neurogenesis and neural cell fate commitment 46. Therefore, we examined mitochondrial content and function in control and NFIA-deleted motor neurons. We first determined if overall levels of ATP were affected in the absence of NFIA. We found that less ATP was produced in tissue collected from E13.5 NFIA-deficient spinal cords (Figure 7B).

**Figure 7.**
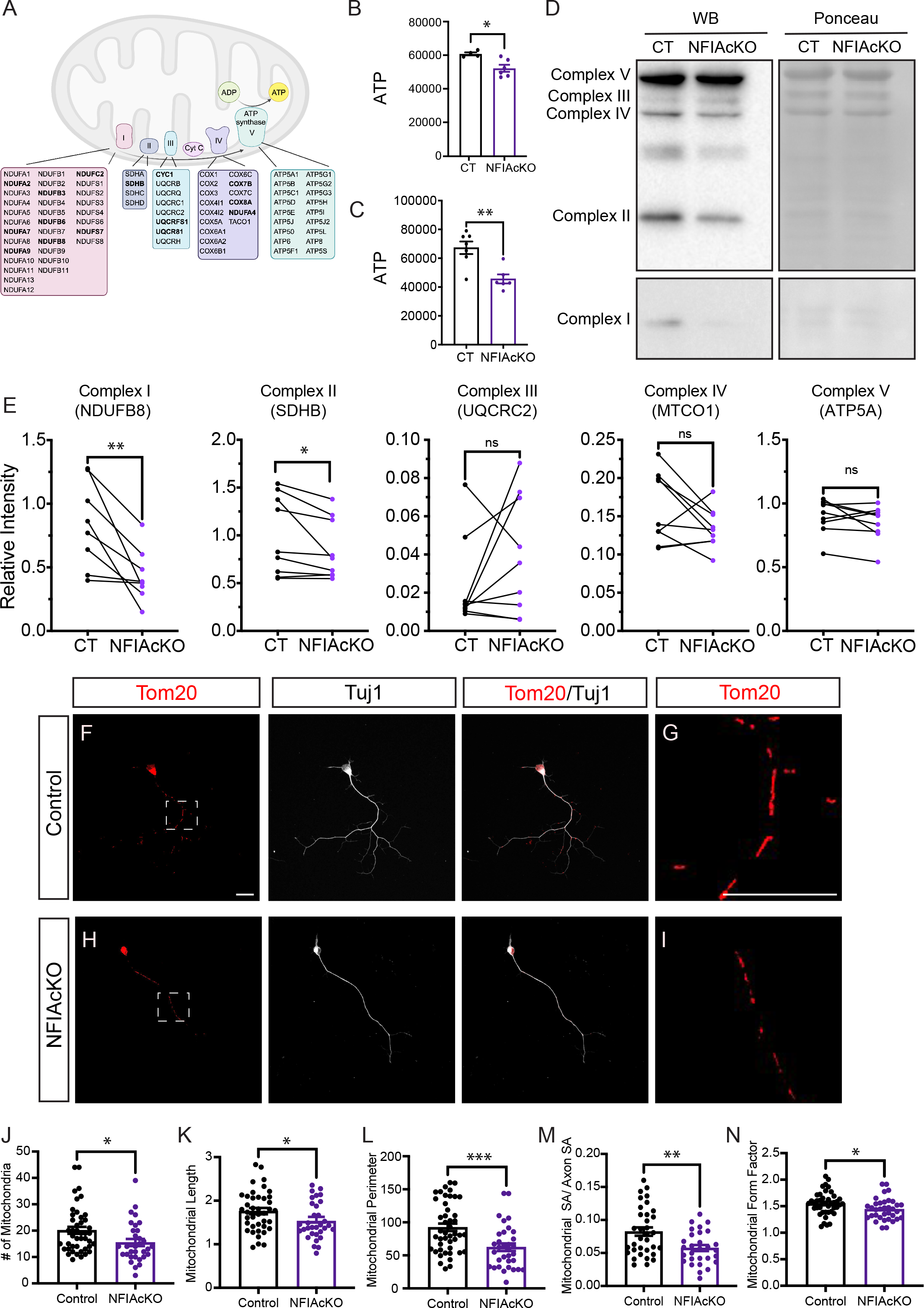
Loss of NFIA leads to mitochondrial dysfunction in motor neurons (A) Schematic representation of the mitochondrial electron transport chain (ETC) and genes associated with complexes I-V. NFIA-bound genes that were differentially expressed in NFIAcKO spinal cords are shown in bold. Figure made with Biorender. (B) Quantification of ATP levels (in Relative Light Units, RLUs) in E13.5 whole spinal cord samples from control (CT) and conditional knockout (NFIAcKO) embryos. (n=5). (C) Quantification of ATP levels in E13.5 ventral spinal cord samples from CT and NFIAcKO embryos. (n=6). (D) Representative image of Western blot (WB) analysis of ETC complexes I-V from CT and NFIAcKO ventral spinal cords. For clarity of the results, the representative western blot for Complex I is shown at a higher exposure than the other complexes. Ponceau staining was used for loading controls and normalization. (E) Quantification of Western blot analysis of mitochondrial OXPHOS complexes. All values are normalized to total protein abundance. (n=7-8). (F,H) Representative images of mitochondria in CT and NFIAcKO motor neuron primary cultures established from E13.5 embryos. Tom20 is used as a mitochondrial marker and Tuj1 as a neuronal marker. (G,I) Higher magnification images of mitochondria (Tom20) from boxed regions in (F,H). (J-N) Quantification of mitochondrial morphology. Motor neuron images were analyzed and quantified using the Mitochondria Analyzer Plugin in Fiji (ImageJ). The measurements for mitochondrial number (J), length (K), perimeter (L), normalized surface area (M), and form factor (N) are shown. Data was obtained from a total of 29-45 motor neurons from 3-4 embryos. All data are represented as mean ± SEM. p* ≤0.05; p** ≤0.01, p***≤0.001 by Student’s T-test. Scale bars represent 20 μm.

Moreover, in the ventral spinal cordーwhere motor neurons are locatedーa marked reduction in ATP production was also observed (Figure 7C), suggesting reduced mitochondrial function in the absence of NFIA. We next assessed the protein levels of oxidative phosphorylation (OXPHOS) genes. From ventral spinal cord lysates, we resolved proteins on SDS-PAGE gel and probed blots with anti-OxPhos antibodies to determine abundance of electron transport chain complex I-IV subunits in control and NFIAcKO motor neurons. This analysis revealed that protein expression of complexes I and II were consistently down-regulated in the absence of NFIA (Figure 7D-E).

Mitochondria change their morphology, size, numbers, and localization throughout development to match the energy demands on the cell ^47,48^. Mitochondria in terminally differentiated neurons have an elongated morphology while mitochondria in immature neurons are more fragmented (Son & Han, 2018). To determine if mitochondrial number and morphology are altered in NFIAcKO motor axons we generated primary motor neuron cultures from E13.5 spinal cords. In primary motor neuron cultures individual control neurons showed elongated mitochondria, while NFIAcKO neurons had fewer, smaller, and shorter mitochondria in their axons, indicative of fragmentation (Figure 7F-I). Quantification of this phenotype in axons revealed that the overall number of mitochondria are reduced in the absence of NFIA (Figure 7J). Further, mitochondrial length (Figure 7K), perimeter (Figure 7L), and surface area (Figure 7M) were also reduced in NFIAcKO motor neurons compared to control neurons. Mitochondrial form factor, which is a measure of shape, also decreased in NFIAcKO motor neurons (Figure 7N). Taken together, these results provide evidence that NFIAcKO motor neurons have reduced mitochondrial content and morphological alterations. These characteristics are consistent with mitochondrial dysfunction and support NFIA as a critical transcriptional activator of motor neuron metabolism.

## DISCUSSION

This study reveals that NFIA directly represses several biological processes crucial for motor neuron development, including motor pool segregation, axonal branching, and NMJ formation. Furthermore, NFIA promotes the transcription of OXPHOS genes, facilitating mitochondrial maturation and marking the first instance of a nuclear transcription factor regulating mitochondrial properties in developing motor neurons. By acting as both a transcriptional activator and repressor, NFIA fine-tunes gene expression to precisely guide motor axons to their skeletal muscle targets during development (Figure 8). These findings establish NFIA as a multifunctional regulator of motor neuron development through a combination of distinct and interconnected molecular mechanisms.

**Figure 8.**
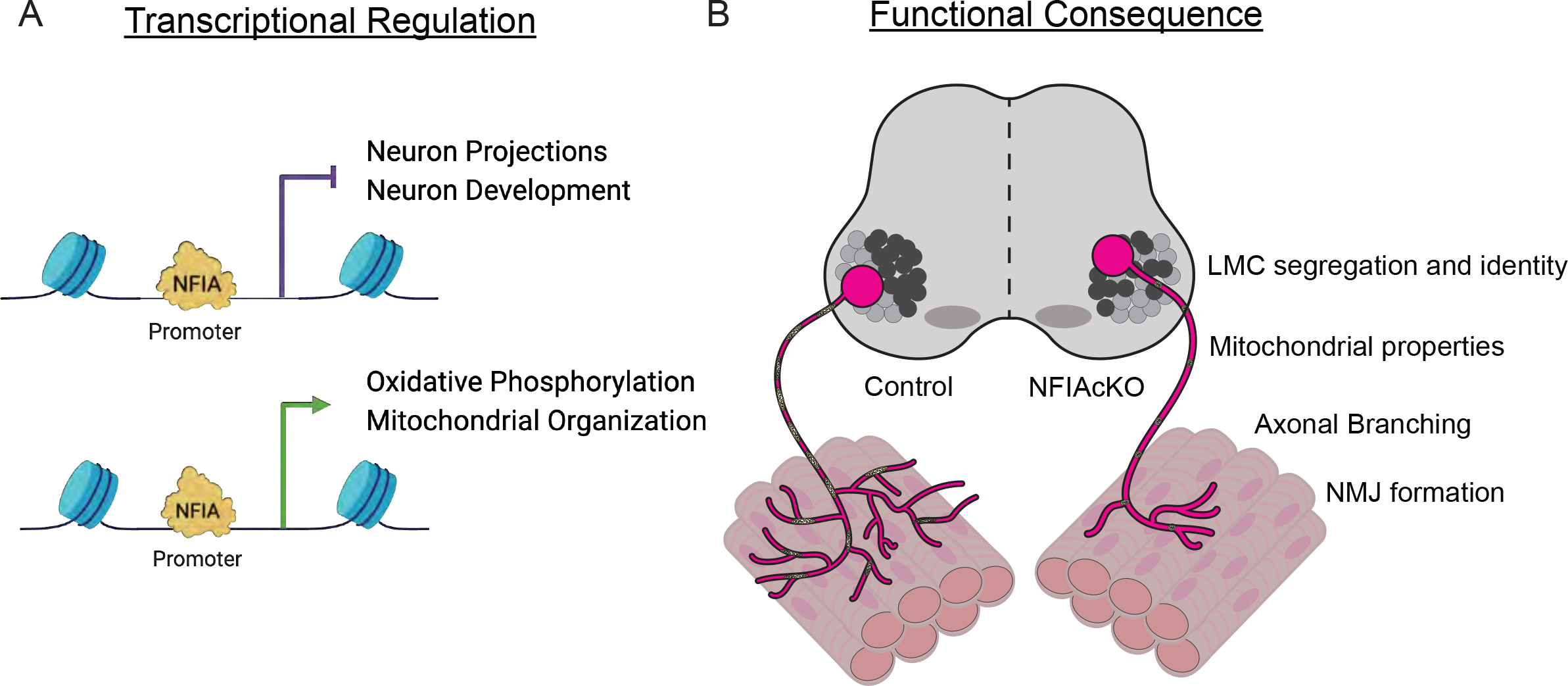
Summary of NFIA function in developing motor neurons (A) Schematic representation of NFIA regulation of gene expression in developing motor neurons. NFIA acts as a transcriptional activator activating genes involved in OXPHOS and mitochondrial properties. NFIA also acts as a transcriptional repressor to affect expression of genes associated with neuron axonal projections and development. (B) Summary of processes affected by loss of NFIA expression (NFIAcKO). When NFIA is deleted, lateral motor columns (LMCs) lack proper segregation; fewer and more fragmented mitochondria are found in motor neuron axons, resulting in less ATP production; axon branching is less complex; and fewer neuromuscular junctions (NMJ) are formed. Figure made in part with Biorender.

### NFIA directly regulates expression of motor neuron genes

Significant progress has been made in our understanding of the initial specification of motor neurons, but the regulatory mechanisms governing the maturation of postmitotic neurons are less clear. Recent studies have started to employ big data analyses to gain a broad view of the molecular landscape of motor neuron development *in vivo* and have shown a surprising amount of molecular diversity in both developing and mature motor neurons ^32,37,50–52^. This includes identification of unique cell-intrinsic transcriptional programs that are initiated in progenitor cells versus post-mitotic cells *in utero,* or exhibit dynamic regulation at postnatal stages ^37,50,51^.

Chromatin accessibility profiling across motor neuron development has revealed at least five transcription factor motifs that are expressed in a temporally dynamic fashion ^37^. One of these motifs belongs to the NFI family of transcription factors, (NFIA, NFIB, NFIC and NFIX) which suggests that these factors may regulate different subsets of genes across motor neuron maturation ^37^.

We started our investigation of NFIA function by combining RNA-seq, ChIP-seq, and ATAC-seq analyses. We identified that NFIA directly regulates >750 genes in E13.5 motor neurons and acts as both a transcriptional activator and repressor. Previous studies on P7 cerebellum and E16 cortex found that NFIA regulated neuron development and neuron projection development genes ^36,53^, indicating that it is important in these processes across the CNS. However, in the brain, NFIA acted as an activator of these genes ^36,53^, whereas in motor neurons it suppressed them. Studies of other transcription factor families, such as Islet, Hox, and Sox, in developing motor neurons have found that they mostly activate expression of neuronal development and axonal guidance genes ^51,54–57^. Conversely, we found that NFIA represses neuron/axon development genes. It is possible that NFIA binds promoters of these genes and prevents other transcription factors from activating them, seeing as there are transcription factor binding motifs near NFIA binding sites.

NFIA also appears to be singular among characterized motor neuron transcription factors in regulating OXPHOS and mitochondrial genes. This suggests that energy metabolism is not ubiquitously controlled by all motor neuron transcription factors, but may be temporally regulated during development. Interestingly, deletion of NFIA in postnatal neural STEM cells, E16 neocortex, and P7 cerebellum did not alter expression of mitochondria-related genes ^36,53,58^, supporting the idea that NFIA regulation of metabolism may be cell type- or stage-dependent.

NFIA has been found to regulate chromatin accessibility in the CNS with other co-factors like Foxp2 ^59^ and NFIB/NFIX ^60,61^. With the exception of regulating chromatin accessibility in human iPSC-derived astrocytes ^62^, compelling evidence of NFIA regulating chromatin accessibility in the CNS (without co-factors) is lacking. In spinal motor neurons, NFIA regulated gene expression without regulating chromatin accessibility, indicating that its regulation of gene expression was not reliant on altered chromatin state.

### Postmitotic expression of NFIA regulates organization of spinal motor neurons

We found that NFIA was expressed in all motor columns and pools except the PGC. PGC is the only motor column that does not innervate skeletal muscle, instead innervating sympathetic chain ganglia ^10^. Therefore, it is possible that regulation of downstream effector pathways in the autonomic system differs from the somatic motor system . Unlike somatic motor neurons, preganglionic motor axons projections were found to not be intrinsically specified ^63^. This suggests that the developmental strategies for forming neuronal connections differs between these two systems.

We found that the absence of NFIA does not alter the generation of limb-innervating motor neurons; however, clustering of LMCM and LMCL motor neurons was disorganized. These cells did not lose their LMC identity, as the number of Isl1/FoxP1 co-positive cells did not change. Instead, they appear to have “confused” LMC identity, co-expressing Isl1/ Lhx1 or Isl1/Hb9. Typically, Lhx1 and Isl1 repress each other’s expression in LMC neurons, which affects the settling of motor neurons into medial and lateral positions ^16,64^. In a subset of NFIAcKO motor neurons, this repression may not occur, or may take place more slowly, resulting in confused identity. We also observed a more dramatic change in dysregulation of motor pool markers, compared to motor column markers. This may be due to the fact that NFIA expression is initiated following motor column specification. Therefore, it is unlikely that NFIA is sufficient for driving early motor neuron column identity, but appears to affect motor pool organization and motor neuron maturation.

In addition to cells with mixed identity, LMCM and LMCL neurons were intermingled in NFIAcKO spinal cords. Netrin/Dcc, Cdh/Catenin, Nectin/Afadin, and Eph/ephrin have previously been shown to regulate motor pool organization ^14,15,17–19,41,65–67^. We found that these systems were dysregulated in NFIAcKO embryos and one of these candidates, Epha4, was upregulated in NFIA mutants. This indicates that NFIA may be working through these pathways to regulate motor neuron position and LMC segregation. A previous *in vitro* study showed that migration of postmitotic cerebellar granule neurons was inhibited using a dominant repressor of all Nfi genes, with Cdh2 and Ephb1 partially responsible for mediating this defect ^68^. Together, these data highlight that NFIA plays a critical role in regulating motor neuron positioning within the LMC, and may have broader roles in regulating cell migration within the CNS.

### NFIA is necessary for regulating terminal branching of motor axons

Many studies have been conducted on the regulation of LMC motor axon invasion into the limb and their selection of dorsal/ventral trajectories ^64,69–73^. However, compared to our knowledge on axon growth, little is known about the molecular regulation of axon branching in motor neurons. Additionally, NFIA knockout has been shown to affect midline crossing of axons in the cortex, but these effects were attributed to cell-extrinsic factors ^74,75^, and whether it had cell-intrinsic effects on axon guidance was not known.

Here, we show that NFIA is an important, cell-intrinsic regulator of axonal branching and terminal arborization, and loss of NFIA results in impairment of NMJ formation in forelimb and hindlimb. Since NFIA mutants exhibited normal limb invasion/bifurcation, its function may be specific to the regulation of axonal branching and terminal arborization. This is similar to ETS transcription factors, such as Pea3, which are expressed following initial motor neuron specification and regulate terminal arborization without affecting initial selection of muscle nerve trajectory ^5^. Our sequencing data reveals that NFIA does not regulate the expression of pool-specific ETS and Nkx factors, therefore it likely regulates axonal branching through a distinct mechanism. Additionally, unlike ETS factors, which are expressed in discrete motor pools, NFIA is broadly expressed in limb-innervating motor neurons. This indicates that it has the potential to regulate terminal arborization, a late step in the formation of motor axon projections, in a broad range of limb muscles.

NFIA directly suppresses genes implicated in axon guidance, including Plexins, Semaphorins, and Ephrin receptors. One possibility is that NFIA represses the expression of genes important for axon guidance that have the potential to inhibit branching ^76^. For example, in mouse and zebrafish, exposure of neurons to increased semaphorin signaling inhibits the formation of new extensions and destabilizes branches ^77,78^. Therefore, upregulation of axon guidance genes in NFIAcKO spinal cords may, in part, explain the stunted and aberrant axonal branches visible in the limb.

Our *in vivo* axonal branching phenotype was recapitulated in *ex vivo* and *in vitro* motor neuron cultures. Previously, a dominant negative repressor for all NFI genes was found to impair neurite outgrowth from cerebellar granule neuron cultures, but not from fully dissociated neurons ^68^. We observed a similar phenotype in the present study, where NFIAcKO explants showed decreased neurite outgrowth, but axon length in isolated motor neurons did not differ from controls. This suggests that NFIA regulation of axon outgrowth differs under conditions of homotypic cell contact. Importantly, NFIAcKO primary dissociated motor neurons had fewer branches compared to controls, further supporting the idea that NFIA mediates cell intrinsic mechanisms of axonal branching.

We also examined the formation of the NMJ in forelimb and hindlimb muscles to determine if projection defects persisted throughout development. Due to the postnatal lethality of NFIAcKO, we analyzed skeletal muscles of these mice at E18.5, prior to birth, when NMJ formation is in progress. Early defects in branching do not correct themselves in NFIA-deleted embryos over time, as evidenced by decreased NMJ formation in NFIAcKO embryos. Deletion of an LMCL transcription factor, Isl2, from motor neurons reduced NMJ formation, which led to impaired gait in adult animals ^55^. Our data demonstrate that embryonic NFIA deletion has persistent consequences and may lead to similar impaired motor output in adults. Overall, NFIA is an important regulator of axon branching and terminal arborization in developing mice, as it represses axon guidance genes that may interfere with this process.

### NFIA regulates mitochondrial size, shape, and respiration in developing motor neuron

During development, neurons adjust their energy balance to meet the high demands of robust axonal growth and branching. Developing neurons undergo a metabolic shift from glycolysis to oxidative phosphorylation (OXPHOS), with an accompanying change in mitochondrial morphology ^45,49^. These highly dynamic organelles are localized to sub-cellular regions with high metabolic demands, such as growth cones, synapses, and axon branches ^29,30,79–82^. The vast majority of proteins involved in mitochondrial structure and function are encoded by nuclear genes, which need to be regulated by transcription factors ^83^. Despite our extensive understanding of mitochondria function in neurons, the precise role of transcription factors in regulating mitochondria function during neuronal development is less clear.

Our study revealed that NFIA, a transcription factor critical for spinal cord development, governs mitochondrial morphology and OXPHOS. NFIAcKO motor neurons displayed reduced levels of OXPHOS molecular components and ATP compared to controls. Mitochondria that do not actively respire are unable to fuse ^83^, and indeed we observed smaller, more fragmented mitochondria in NFIA-deficient motor neurons. The observed defects in axonal branching and NMJ formation in NFIAcKO motor neurons suggests that regulation of ATP production is a vital aspect of motor neuron maturation and NFIA function. Apart from directly regulating expression of axon guidance molecules/receptors, NFIA also regulates mitochondrial respiration, providing neurons with the energy needed to establish connections with their targets. Transcription factors such as Tfam, Nrf1, NeuroD6, and Foxg1 have previously been linked to mitochondrial function during neurodevelopment ^84–89^. However, of the transcription factors known to regulate motor neuron development (e.g. Hox, Fox, LIM, ETS), NFIA is the only one, to our knowledge, that has been shown to regulate mitochondrial properties. Our data, combined with recent evidence that NFIA regulates expression of OXPHOS genes in the adipose tissue ^90^, affirms that NFIA is important for regulating metabolism across tissue types both during development and in adult tissues.

Outside of OXPHOS, our sequencing data indicates that NFIA regulates other aspects of mitochondrial functions, including mitochondrial morphology and transport. Future studies examining processes such as mitochondrial motility would provide insight into the breadth of mitochondrial regulation by NFIA. Altogether, our study establishes NFIA as an important regulator of mitochondrial function in neuronal development.

## CONCLUSIONS

Broad multi-omic profiling of transcription factors important in development can not only help us correlate transcriptional programs with phenotypes, but also define unknown functions of these transcription factors. NFIA has long been considered a glial-specification factor; however, our findings demonstrate its critical roles in the development of spinal motor neurons by directly regulating multiple molecular processes. NFIA emerges as a promising transcription factor for future studies aimed at gaining a comprehensive understanding of terminal arborization and mitochondria function in developing neurons. The observed impaired neurite outgrowth from spinal cord explants and branching from isolated motor neurons in NFIAcKO mice has implications for axonal regeneration. Investigating the connection between NFIA, mitochondrial metabolism, and regeneration presents an intriguing avenue for future research. Given the association between mitochondrial properties and neurodevelopmental/neurodegenerative disorders, elucidating NFIA’s role in neuronal development holds promise for shedding light on these pathologies. Understanding the involvement of NFIA in neuronal development may offer insights into the underlying mechanisms of such disorders.

## METHODS

### KEY RESOURCE TABLE

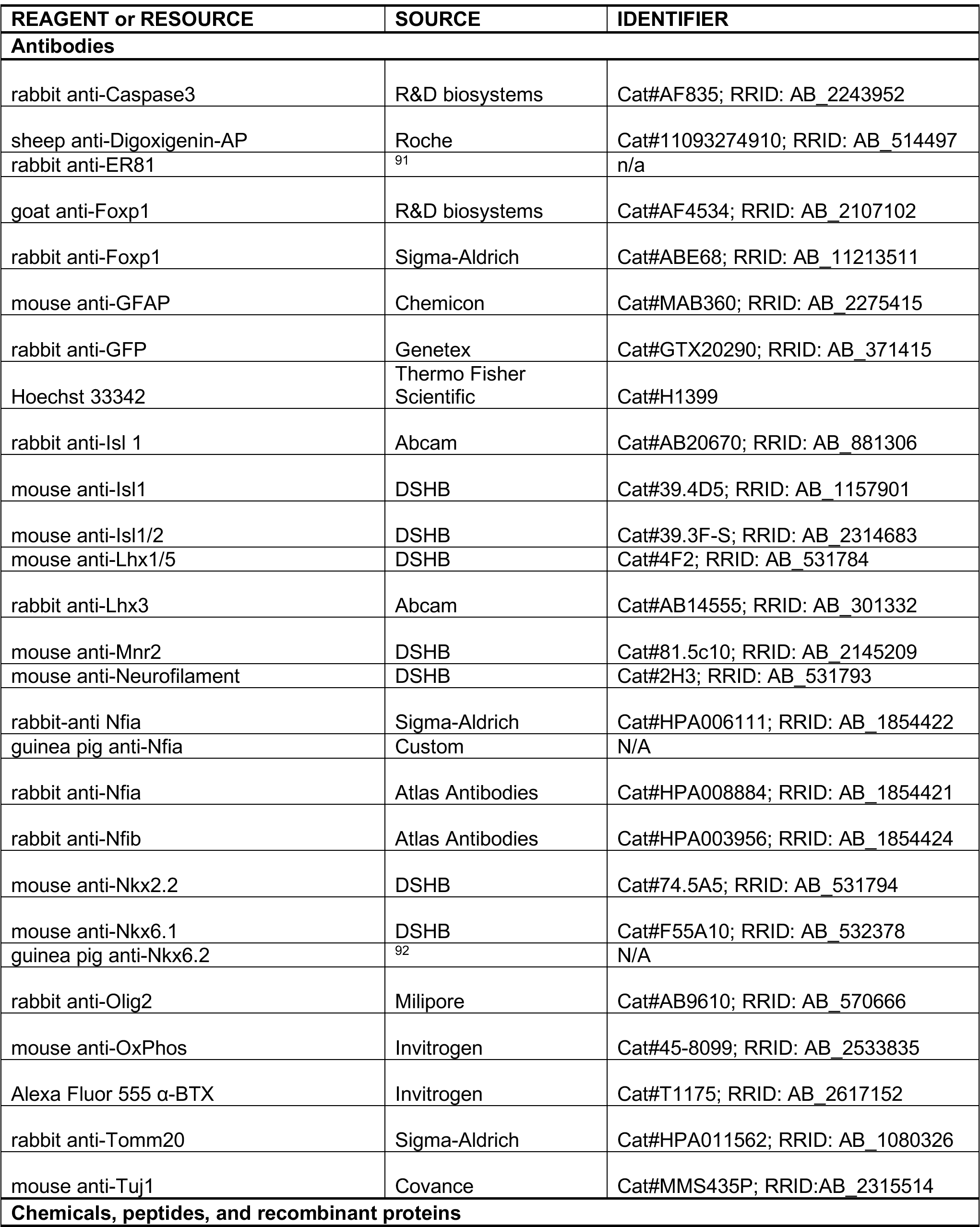
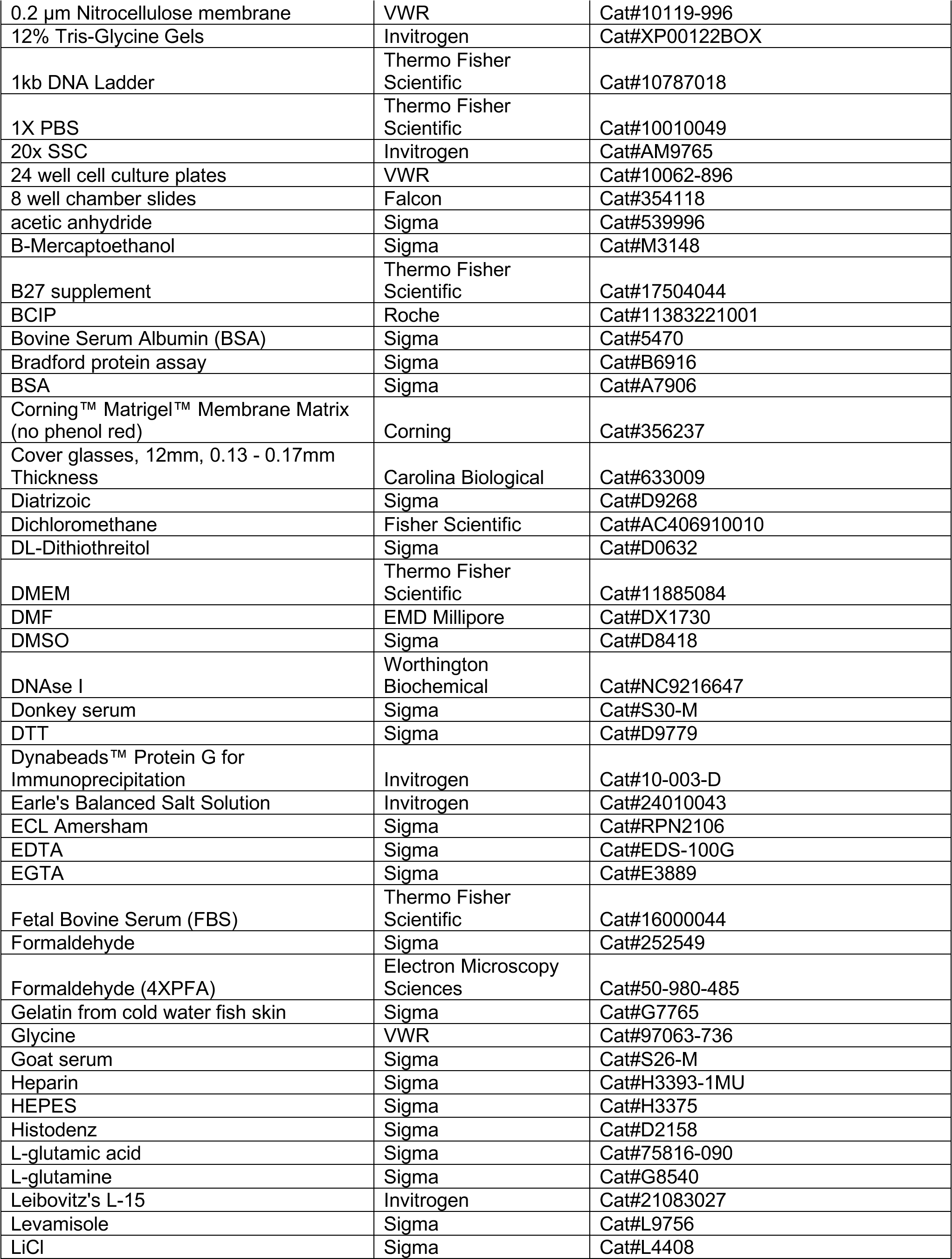
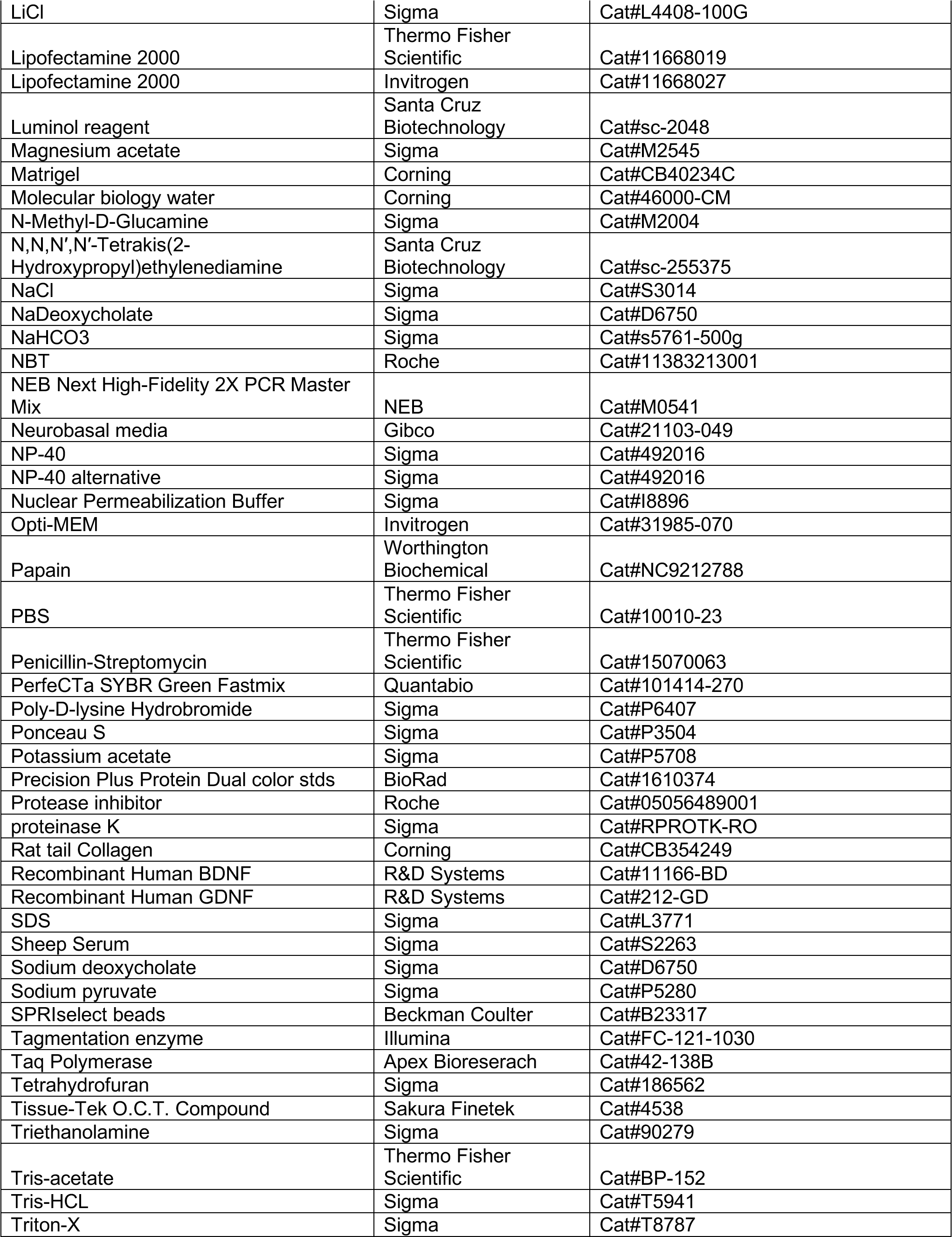
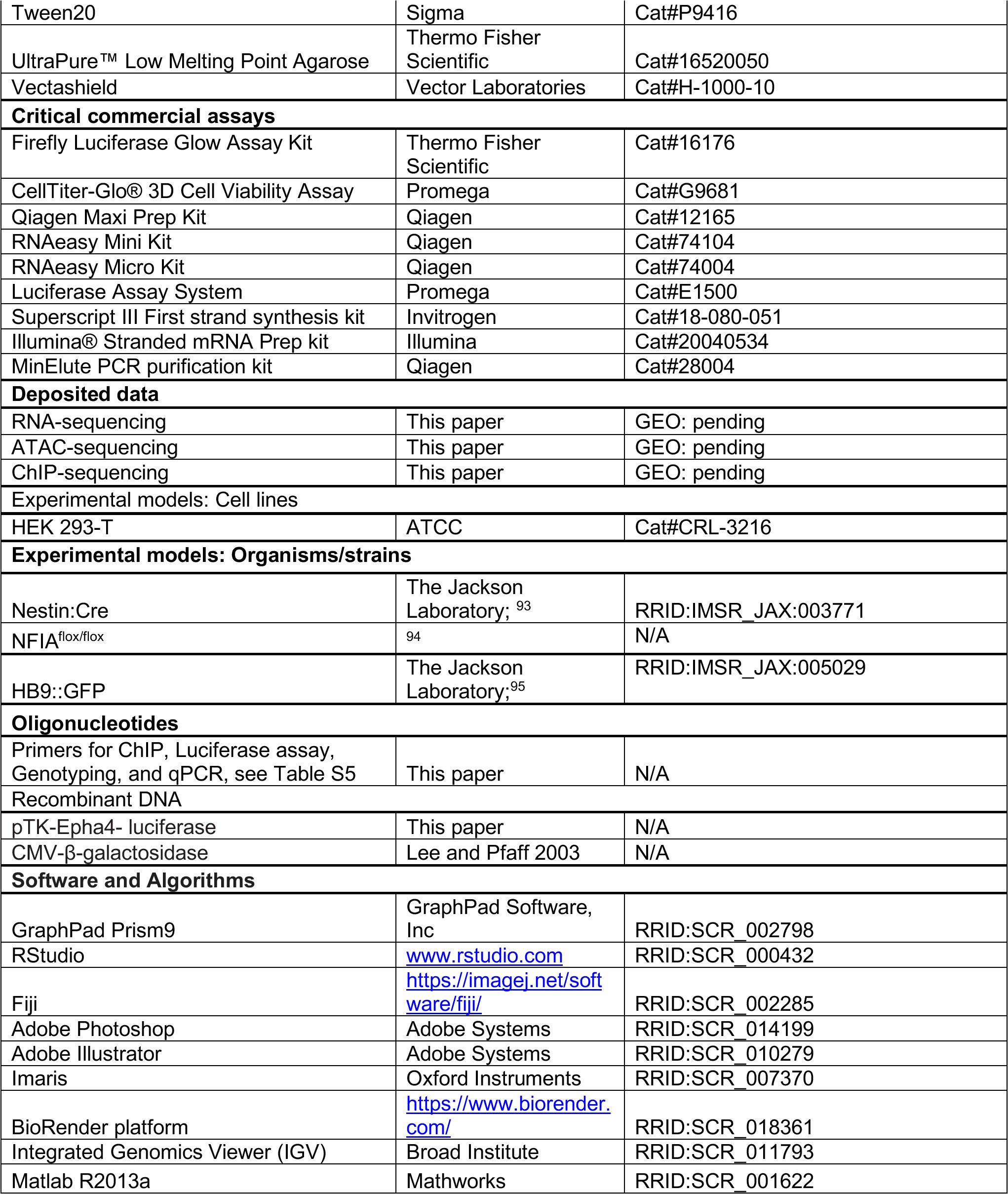

### Animal Models

NFIA conditional knockout mice (NFIAcKO) were generated by crossing NFIA^fl/fl^ ^94^ mice to Nestin-cre (Jackson Laboratory 003771) mice. All mice were maintained on a c57BL/6J background. Both male and female mice were used in all experiments and were randomly divided into experimental groups. HB9-positive cells and corresponding axonal projections were examined using the HB9::GFP (Jackson Laboratory 005029) reporter mouse line. Care of all animals in this study followed NIH guidelines and procedures were approved by the University of California San Diego Institutional Animal Care and Use Committee (IACUC). PCR genotyping was performed on harvest tail clippings that were incubated in 50mM NaOH for 12 min at 95°C. Tail preps were neutralized with 1M TRIS pH8 and 5mM EDTA pH8, then used for PCR genotyping. Primers used for genotyping can be found in Table S5.

### Immunohistochemistry

Embryos were harvested between gestational day E10.5-P4 and fixed in 4% formaldehyde, cryoprotected with 20% sucrose overnight, embedded in O.C.T compound (Tissue-Tek), and cryosectioned at 14µm. Tissues were subjected to either immunohistochemistry or in situ hybridization. For immunohistochemistry, cryosections were washed in 1XPBS with 0.01% tween (PBS-T) for 10 min three times, followed by heat-induced antigen retrieval for 15 min at 95°C in 10 mM sodium citrate pH6 with 0.05% Tween-20. For primary cultures, the heat-induced antigen retrieval step was omitted. After 1h of blocking (10%serum/1% fish gelatin/0.2% Triton X/ 1XPBS), Primary antibodies were applied overnight at 4°C. Slides were then washed three times in 1XPBS-T for 5 min each and appropriate goat or donkey secondary antibodies (Invitrogen) were applied at 1:500 for 1-2 h. Slides were then washed three times in 1XPBS-T for 5 min, followed by Hoechst 33258 staining at 1:8000 for 10 min and then washed once with 1XPBS. Slides were mounted with Vectashield (Vector laboratory) and imaged.

To analyze dorso-ventral axon trajectories in cross section, CT and NFIAcKO E13.5 embryos carrying the Hb9::GFP transgene were fixed in 4%formaldehyde overnight, embedded in 4% low-melting point agarose and sectioned at 400 μm using the McIlwain Tissue Chopper (Stoelting). Sections were incubated with primary antibodies in blocking buffer for 48 h at 4°C. After incubation with secondary antibodies (1:500) for 3h at room temperature, sections were washed and mounted on slides with Vectashield (Vector Laboratories). Sections were imaged on an Olympus BX63 fluorescence microscope.

Primary antibodies used were as follows: rabbit anti-Caspase3 (R&D biosystems; 1:500), rabbit anti-ER81 (Arber et al., 2000; 1:30,000), goat anti-Foxp1 (R&D biosystems; 1:100), rabbit anti-Foxp1 (Sigma-Aldrich; 1:100), rabbit anti-GFP (Genetex, 1:700), rabbit anti-Isl1 (Abcam; 1:400), mouse anti-Isl1 (DSHB; 1:25), mouse anti-Isl1/2 (DSHB; 1:50), mouse anti-Lhx1/5 (DSHB; 1:5), rabbit anti-Lhx3 (Abcam; 1:400), mouse anti-Mnr2 (DSHB; 1:10), mouse anti-Neurofilament (DSHB; 1:70), rabbit-anti Nfia (Sigma-Aldrich; 1:500), rabbit anti-Nfib (Sigma-Aldrich; 1:300), mouse anti-Nkx2.2 (DSHB; 1:5), mouse anti-Nkx6.1 (DSHB; 1:10), guinea pig anti-Nkx6.2 (Vallstedt et al., 2001; 1:700), and rabbit anti-Olig2 (Sigma-Aldrich; 1:250). Custom anti-Nfia antibody was generated in guinea pig following the peptide sequence reported in ^35^(1:300 dilution).

### *In Situ* Hybridization

To detect Eph4a, we generated mRNA probes using specific primers found on the Allen Atlas Database, and performed *in situ* hybridization on mouse spinal cord tissue. *In situ* hybridization on frozen embryos was performed as previously described ^35^. In short, after initial washes with PBS, deproteinization was conducted by proteinase K (10 μg/ml) treatment for 5 min. This was followed by 4% paraformaldehyde for 15 minutes, 0.25% acetic anhydride in 0.1M triethanolamine for 5 min, and washing with 1XPBS for 5 min. A pre-hybridization step at 65°C for 1h was followed by hybridization of the RNA probe at 65°C overnight. Chromogenic detection was done the following day. Sections were first washed with 1X SSC for 15 min at 65°C, then by a higher stringency wash of 0.5X SSC at 65°C for 30 min. This was followed by rinsing with PBT (1xPBS, 2mg/mL BSA, 0.1% Triton X-100), 1h blocking in 20% sheep serum (in PBT), and 2h incubation with sheep anti-Digoxigenin-AP (Roche; 1:2000) at room temperature. Sections were then washed with PBT and incubated for 10 min in AP buffer (100mM Tris, 50mM MgCl2, 100mM NaCl, 0.1%Tween-20, 5mM levamisole). The sections were developed in NBT/BCIP in the dark overnight. The reaction was stopped with AP buffer once the signal was developed and treated with 4%PFA for 10 min. Slides were then mounted with glycerol.

### Image analysis

Motor neurons populating motor column and motor pool divisions were identified by transcription factor expression in 14 μm sections of E13.5 spinal cord. To determine the number of motor neurons present at brachial and lumbar levels, as well as, cells that co-express motor column/pool markers, cells were counted unilaterally using ImageJ 96 in ≥3 embryos of each genotype. To create contour plots, motor neuron positions were acquired using the ‘‘spots’’ function using Imaris software (Bitplane) to assign x and y coordinates. Coordinates were expressed relative to the spinal cord midline, defined as positions x = 0, y = 0. Height was defined as the distance from the ventral limit of the central canal to the dorsal-most edge of the spinal cord, and hemichord width as the distance from the central canal to the most lateral edge. Relative dorso-ventral and medio-lateral positions of motor neurons were expressed as percentages of spinal cord height and hemicord width, respectively. Dorsoventral and mediolateral positions were plotted using Matlab software R2013a (Mathworks) using the ksdensity function. Analysis of EphA4 integrated density was performed on thresholded spinal cord images using ImageJ in ≥4 embryos of each genotype.

### RNA-sequencing

RNA was harvested from control and NFIAcKO E13.5 mouse spinal cords (n=3) using the Qiagen RNeasy Mini kit was used for library preparation. Total RNA quality was assessed using an Agilent Tapestation 4200, and samples with an RNA Integrity Number (RIN) >8 were used to generate RNA sequencing libraries using the Illumina® Stranded mRNA Prep kit (Illumina).

Resulting libraries were multiplexed and sequenced with 100 basepair (bp) paired-end reads (PE100) to a depth of approximately 25 million reads per sample on the Illumina NovaSeq 6000 platform at the UCSD IGM Genomic Center. Samples were demultiplexed using bcl2fastq Conversion Software (Illumina). Adaptor trimmed FASTQ paired- end reads were aligned using STAR (v.2.6.1a) to a mouse reference genome (mm10; --genome Dir /mouse/mm10/ -- sjdbGTFfile gencode.vM10.primary_assembly.annotation.gtf). Transcripts were quantified using HTSeq, data was normalized using RUV-seq and differential gene expression analysis was conducted using DESeq2 ^97^. Significant differentially expressed genes (DEGs) had an FDR < 0.05, log2fold change ≥ 0.3 or ≤0.3. Gene ontology (GO) analysis was performed using the ToppGene Suite^98^. BigWig files were generated using deeptools, bamCoverage --bam {} -- binSize 20 --smoothLength 60 --normalizeUsing CPM.

### Real-time Quantitative PCR (qPCR)

RNA was harvested from E13.5 mouse spinal cords using the Qiagen RNeasy Mini kit and complementary DNA (cDNA) libraries were prepared from each RNA isolate with the SuperScript III reverse transcriptase kit (ThermoFisher Scientific) and an anchored oligo-dT18 primer. Gene-specific primers for mouse (Key resources table) were used for quantitative PCR using the PerfeCTa SYBR Green Fastmix (Quantabio). To ensure that single amplicons were produced in each PCR reaction, PCR products were separated on 2% agarose gels, and melting curve protocols were performed after each qPCR run. Real-time PCR fluorescence measurement and melt curve analyses were performed using a Bio-Rad C1000^TM^ Thermal Cycler equipped with a CFX384^TM^ System (Bio-Rad). Transcript expression levels were quantified and normalized relative to Cyclophilin (Cyc) using the ΔΔ cycle threshold (ΔΔCT) method. The PCR program used was as follows: 30 s at 95°C followed by 40 cycles of 5 s at 95°C/30 s at 60°C, and 12 s at 72°C. Normalized abundance of all NFIAcKO transcripts was standardized to CT transcripts, which were set to 100%. Student’s t-test was also performed to confirm that transcript abundance of Cyclophilin did not significantly differ between tissues. Primers used for qPCR analysis can be found in Table S5.

### ChIP-sequencing

After harvesting, samples were flash frozen and sent to Active Motif (Carlsbad, CA) for ChIP-Sequencing. Active Motif prepared chromatin, performed ChIP reactions, generated libraries, sequenced the libraries and performed basic data analysis. In brief, wildtype mouse E13.5 spinal cords were homogenized and fixed with 1% formaldehyde for 15 min and quenched with 0.125 M glycine. Chromatin was isolated by adding lysis buffer, followed by disruption with a Dounce homogenizer. Lysates were sonicated and the DNA sheared to an average length of 300-500 bp with EpiShear Probe Sonicator (Active Motif). Genomic DNA (Input) was prepared by treating aliquots of chromatin with RNase, proteinase K and heat for de-crosslinking, followed by SPRI beads clean up (Beckman Coulter) and quantitation by Clariostar (BMG Labtech). Extrapolation to the original chromatin volume allowed determination of the total chromatin yield. An aliquot of chromatin (40 μg) was precleared with Protein G beads (Invitrogen). Genomic DNA regions of interest were isolated using 30 μL of rabbit anti-NFIA antibody (Atlas Antibodies). Complexes were washed, eluted from the beads with SDS buffer, and subjected to RNase and proteinase K treatment. Crosslinks were reversed by incubation overnight at 65°C, and ChIP DNA was purified by phenol-chloroform extraction and ethanol precipitation.

Illumina sequencing libraries (a custom type, using the same paired read adapter oligonucleotides described by ^99^ were prepared from the ChIP and Input DNAs on an automated system (Apollo 342, Wafergen Biosystems/Takara). After a final PCR amplification step, the resulting DNA libraries were quantified and sequenced on Illumina’s NovaSeq (75 bp reads, single end). Reads were aligned to the mouse genome (mm10) using the BWA algorithm (default settings; Li & Durbin, 2009). Duplicate reads were removed and only uniquely mapped reads (mapping quality >= 25) were used for further analysis. Alignments were extended in silico at their 3’-ends to a length of 200 bp, which is the average genomic fragment length in the size-selected library, and assigned to 32-nt bins along the genome. The resulting histograms (genomic “signal maps”) were stored in bigWig files. Peak locations were determined using the MACS algorithm (v2.1.0; ^101^) with a cutoff of p-value = 1e-7. Peaks that were on the ENCODE blacklist of known false ChIP-Seq peaks were removed. Signal maps and peak locations were used as input data to Active Motifs proprietary analysis program, which creates Excel tables containing detailed information on sample comparison, peak metrics, peak locations and gene annotations.

### ATAC-seq

NFIA^ff^;Hb9::GFP and NFIA^ff^;Nestin-cre;Hb9::GFP E13.5 mouse spinal cords (n=2) were dissected in ice-cold PBS. Cells from each spinal cord were dissociated with 500 μL Papain (10U; Worthington Biochemical) and DNAse I (100 U; Worthington Biochemical) in Earle’s Balanced Salt Solution (Gibco) for 30 min at 37°C. Cells were washed and transferred to FACS buffer (0.1% BSA, 1% Pen/Strep, 10 mM HEPES, 5U/mL DNase I powder in Leibowitz medium). Cells were analyzed and sorted using a BD FACS Aria II Cell Sorter (UCSD Human Embryonic Stem Cell Core Facility), and GFP+ cells were isolated. Freshly-sorted cells were resuspended in cryopreservation solution (50% FBS/40% DMEM^H^, 10% DMSO) and frozen at- 80°C.

Permeabilized nuclei were obtained by resuspending cells in 250µL Nuclear Permeabilization Buffer (0.2% IGEPAL-CA630, Sigma), 1mM DTT (Sigma), Protease inhibitor (Roche), 5% BSA (Sigma) in PBS (Thermo Fisher Scientific)], and incubating for 5 min on a rotator at 4°C. Nuclei were then pelleted by centrifugation for 5 min at 500xg at 4°C. The pellet was resuspended in 25µL ice-cold Tagmentation Buffer [33 mM Tris-acetate (pH = 7.8) (Thermo Fisher Scientific), 66 mM K-acetate (Sigma), 11 mM Mg-acetate (Sigma), 16 % DMF (EMD Millipore) in Molecular biology water (Corning)]. An aliquot was then taken and counted by hemocytometer to determine nuclei concentration. Approximately 50,000 nuclei were resuspended in 20µL ice-cold Tagmentation Buffer, and incubated with 1µL Tagmentation enzyme (Illumina) at 37 °C for 60 min with shaking 500 rpm. The tagmentated DNA was purified using MinElute PCR purification kit (Qiagen). The libraries were amplified using NEB Next High-Fidelity 2X PCR Master Mix (NEB) with primer extension at 72°C for 5 min, denaturation at 98°C for 30s, followed by 8 cycles of denaturation at 98°C for 10 s, annealing at 63°C for 30 s and extension at 72°C for 60 s. Amplified libraries were then purified using MinElute PCR purification kit (Qiagen), and two size selection steps were performed using SPRIselect beads (Beckman Coulter) at 0.55X and 1.5X bead-to-sample volume ratios, respectively. QC data processing was performed as described in ^102^.

Paired-end FASTQ reads for each replicate were aligned using Bowtie 2.0 (bowtie2 – very-sensitive -k 30) to the mouse genome (mm10). Samtools was used to sort (samtools sort), remove duplicates (samtools markdup), remove mitochondrial reads (samtools view) and index BAM files (samtools index). BAM files were shifted by +4bp and -5bp using an in-house algorithm ^100^. Differential accessibility of promoters analysis between CT and NFIAcKO samples was performed using Diffbind v1.16.3 ^103^. To determine whether accessible chromatin was enriched for NFIA binding sites on promoters (-2000bp, +500bp TSS), a randomized permutation test was performed in R using scripts described in ^104^. To determine the motifs enriched near NFIA binding sites, HOMER motif analysis was performed. ATAC-seq regions were split into peaks occurring within promoters and the GimmeMotifs function ^105^ was used with the Homer ^39^ motif-finding algorithm.

### Chromatin Immunoprecipitation (ChIP)

Mouse E13.5 spinal cords were dissected, dissociated, and processed for ChIP assays. Five E13.5 mouse dissected spinal cords were used for each immunoprecipitation and control IgG reaction in this assay. Harvested cells were washed in 1XPBS buffer then fixed with 1% formaldehyde for 10 min. Cells were washed first with buffer I (10 mM EDTA, 0.5 mM EGTA, 0.25% TritonX100, 10 mM HEPES pH6.5), then with buffer II (0.2M NaCL, 1 mM EDTA, 0.5 mM EGTA, 10 mM HEPES pH6.5). Cells were resuspended in lysis buffer (0.5% SDS, 5 mM EDTA, 25 mM Tris-HCl pH8.0, protease inhibitors), and sonicated 10 times for 10s each using a Branson Sonifier 450 Sonicator. Supernatants were diluted five times in dilution buffer (2mM EDTA, 150mM NaCl, 1% TritonX100, 20mM Tris-HCl pH8.0, protease inhibitors). The diluted samples underwent an immunoclearing step with protein G agarose beads (Santa Cruz Biotechnology). Cell lysates were subject to immunoprecipitation using rabbit anti-NFIA antibody (Sigma) and protein G agarose beads (control) overnight. The next day, beads were subjected to 10 min washing steps with TSE I buffer (0.1% SDS, 1% TritonX100, 2 mM EDTA, 150 mM NaCl, 20 mM Tris-HCl pH8.0), TSE II buffer (0.1% SDS, 1% TritonX100, 2 mM EDTA, 500 mM NaCl, 20 mM Tris-HCl pH8.0), buffer III (0.25M LiCl, 1% NP-40, 1% deocycholate, 1 mM EDTA, 10 mM Tris-HCl pH8.0) and lastly 1XTE buffer. Genomic DNA fragments were eluted with elution buffer (1% SDS, 0.1M NaHCO3), and cross-linkage was reversed by incubating at 65°C overnight. Samples were treated with proteinase K (Promega) at 42°C for 2h. The genomic DNA fragments were purified using phenol/chloroform extraction and the DNA pellet was resuspended in water. For ChIP-qPCR, ChIP DNA was analyzed by quantitative PCR (BioRad) using PerfeCta SYBR Green Fastmix (Quanta Biosciences) according to manufacturer’s instructions. The PCR program used was as follows: 2 min at 95°C followed by 45 cycles at 10s at 95°C /45s at 57°C and 12s at 72°C. PCR was performed using region specific primers listed in Table S5. All ChIP experiments were performed in triplicate and at least two independent times.

### Luciferase Reporter assay

Genomic DNA was purified and PCR was performed using region-specific primers (Table S5). Amplified regulatory regions were cloned into pTK-Luciferase reporter plasmid. HEK293T cell lines cells were transfected with pTK-luciferase constructs and a CMV-β-galactosidase vector using Lipofectamine 2000 (Thermo Fisher Scientific). Cells were harvested and analyzed for luciferase activity using the Luciferase Assay System (Promega) according to manufacturer instructions. Luminescence was read on a Victor X3 MultiLabel Plate Reader (Perkin Elmer). β- galactosidase was used to normalize for transfection efficiency.

### Clearing and whole-mount immunostaining

Embryo clearing was modified from previously described protocols ^106,107^. In short, E13.5 CT and NFIAcKO embryos carrying the Hb9::GFP transgene were fixed overnight in 4% formaldehyde in PBS. Embryos were decolored in 25% N,N,N′,N′-Tetrakis(2-Hydroxypropyl)ethylenediamine (Santa Cruz Biotechnology) in PBS for 2 d at 37°C. This was followed by dehydration in a tetrahydrofuran (Sigma) series (1 h in 50%, 70%, 90% THF in sterile water at 4°C), 1h incubation in dichloromethane (Fisher) at 4°C, and rehydration into sterile water. After 4 washes in water, samples were incubated in glycine solution (0.2% TritonX-100/20% DMSO/300 mM glycine/1XPBS) overnight at 37°C. Following this, samples were blocked (0.2% TritonX-100/10% DMSO/6% donkey serum/1XPBS), washed in 0.2% PBS-T with 10 μg/mL heparin (PTwH) for 2h. Primary rabbit anti-GFP antibody (1:150; Genentex) was diluted in primary antibody buffer (5% DMSO/3% donkey serum/PTwH) and incubated for 3 days at 37°C. Samples were washed overnight in PTwH at room temperature, and then incubated with goat anti-rabbit Alexa Fluor 488 secondary antibody (1:500; Thermo Fisher Scientific) diluted in secondary antibody buffer (3% donkey serum/PTwH) for 3 days at 37°C. After washing overnight at room temperature in PTwH, samples were incubated in refractive index matching solution (53.2g Histodenz, 0.6g Diatrizoic acid, 1g NMDG added to 20 mL sterile water; RI=1.553) overnight at 37°C and imaged using a Zeiss Z.1 light sheet microscope.

### Analysis of NMJ

For NMJ staining, E18.5 embryos were fixed in 4% PFA overnight at 4°C then bicep and gastrocnemius muscles were harvested. Samples were incubated in 0.1M glycine in 1XPBS, then immunostained with mouse anti-neurofilament (DSHB; 1:70) overnight at 4°C, followed by secondary antibody and Alexa Fluor 555 α-BTX (Invitrogen; 1:500) for 2 h at room temperature. NMJ images were acquired on a Nikon AX R confocal microscope. To analyze NMJs, maximum intensity projections were created using ImageJ software. The number of NMJs was quantified along the entire nerve within muscles. Quantification of the number of secondary branches and NMJ (signified by co-localization of neurofilament and α-BTX) was performed using ImageJ software.

### Explant and primary motor neuron cultures

Motor neurons were cultured using modified, previously described protocols ^55,108^. Briefly, CT and NFIAcKO E13.5 mouse ventral spinal cords were dissected in cold PSB using ultra-fine scalpels and maintained in Plain Neurobasal Media (NBM; Gibco; 21103049) for further processing. For explants, embryos carrying the Hb9::GFP transgene were used. Using 8-well chamber glass slides (Falcon), explants were embedded in roughly 60μL of a collagen-Matrigel (1:1) mixture consisting of Collagen I (Corning), Matrigel (Corning), and 10XPBS solution (60% of 10x PBS, 115 mM NaOH in cell culture grade water) in a 5:5:1 ratio. The explant-gel mixtures were allowed to sit at room temperature for 30 minutes to solidify. For isolated cells, ventral spinal cords were dissociated with 500 μL Papain (10U; Worthington Biochemical) and DNAse I (100 U; Worthington Biochemical) in Earle’s Balanced Salt Solution (Gibco) for 30 min at 37°C. Cells were washed and resuspended in Complete NBM (NBM with 2% B-27, 1% Pen/step, 1% L-glutamic acid, 0.4% L-glutamine). Following, 25,000 cells were plated onto 12 mm glass coverslips (Carolina Biological) coated with poly-D-lysine (>300,000 MW; Sigma). Explants and isolated neurons were incubated in Complete NBM with 100 μg/mL GDNF, 20 μg/mL BDNF (R&D Systems) for 2-4 days, then fixed in 4% formaldehyde for 10 minutes. Explants were immunostained with rabbit anti-GFP (Genentex; 1:700), mouse anti-Tuj1 (Covance; 1:500), and/or rabbit anti-Tomm20 (Sigma; 1:500) antibodies and mounted with Vectashield prior to imaging on a Nikon AX R confocal microscope.

Sholl analysis was performed using the Sholl analysis plugin after thresholding each neuron using Image J, and ≥ 6 explants per condition per experiment were analyzed. For isolated motor neurons, axon length, branches, surface area and mitochondrial length analyses were performed on thresholded images in ImageJ. At least 10 motor neurons per embryo, from 3-4 embryos, were used for analysis. Mitochondrial number, area, perimeter and form factor were analyzed using the Mitochondrial Analyzer ^109^ plugin.

### Luminescent ATP assay

ATP levels in motor neurons were assessed using the bioluminescent reagent Cell Titer-Glo 2.0 (Promega, Madison, WI). CT and NFIAcKO E13.5 mouse spinal cords (whole and ventral) were dissected in cold PSB and dissociated with 500 μL Papain/DNAse I (Worthington Biochemical) in Earle’s Balanced Salt Solution (Gibco) for 30 min at 37°C. Cells were washed and resuspended in NBM (Gibco). 50,000 cells were incubated with CellTiterGlo reagent (Promega) according to manufacturer instructions. Luminescence was read on a Victor X3 MultiLabel Plate Reader (Perkin Elmer).

### Western blot analysis

Open book preparations of E13.5 mouse spinal cords were dissected and ventral spinal cords were separated. Ventral spinal cords were lysed in 200 μL of chilled lysis buffer composed of 50mM Tris-HCl pH 7.5, 0.25M NaCl, 5mM EDTA and EGTA, 1mM dithiothreitol, 0.1% Triton X- 100, 0.1% deoxycholate, 0.1% NP-40, and protease inhibitor (Roche). Protein lysates were quantified using the Bradford protein assay (Thermo Fisher Scientific), and 20 μg were electrophoretically separated on 12% Tris-Glycine Gels (Invitrogen) and transferred onto 0.2 μm nitrocellulose membranes. Membranes were then washed in TBS-T (10 mM Tris-Cl, 150 mM NaCl, 0.05% (v/v) Tween 20, pH 7.4) and blocked for 1h at room temperature in TBS-T containing 5% skimmed milk powder. After blocking, the membranes were incubated overnight at 4°C with mouse monoclonal anti-OxPhos antibody (1:1000). Blots were incubated with goat anti- mouse secondary antibody conjugated to horseradish peroxidase (Jackson immunological; 1:5000 in 5% milk TBS-T) at room temperature for 1h. Membranes were imaged following 1 to 5 min incubation in Luminol reagent (Santa Cruz Biotechnology). Equal protein content among samples was confirmed via Ponceau staining. Quantification of bands observed on Western blots was performed using ImageJ ^96^and standardized to corresponding total protein on lanes of Ponceau-stained gels.

### Statistical methods

All p-values and replicate numbers were described in the figure legends. Data throughout the paper is represented as mean ± SEM unless otherwise noted. Statistical analysis was performed with GraphPad Prism v10 for MacOS. One-way Anova test followed by Tukey’s test was used to determine the differences between group or condition means, with significant differences reported as different uppercase letters. Student’s two-tailed t-test was used to compare individual means and is reported as asterisks in associated figure graphs. Nested t-test was used to determine significance in Sholl analysis. For Western blot analysis, pairwise t-tests were used to compare protein abundance among litters. No statistical methods were used to pre-determine sample sizes, but our sample sizes are similar to those reported in previous publications ^33,35,110^.

## RESOURCE AVAILABILITY

### Lead Contact

Requests for reagents or resources should be directed to the Lead Contact, Stacey Glasgow

### Material availability statement

Reagents generated for this study are available upon request from the Lead Contact.

### Data code availability

Data and code pertaining to this study are available upon request. The accession numbers for RNA-seq, ChIP-seq and ATAC-seq data reported in this paper are: GEO: ####. Lists of genes identified in this study are provided in Table S1-S4.

## AUTHOR CONTRIBUTIONS

J.G and S.M.G conceived the project and designed the experiments. J.G and S.M.G wrote the manuscript. J.G, K.M., K.J., S.A., S.J. and S.M.G performed the experiments. J.G and S.M.G analyzed the data. K.M, A.M.S, and K.J performed data analysis for specific figures. J.G and H.S.M performed bioinformatic analysis of sequencing experiments.

## ACKNOWLEDGEMENTS

We gratefully acknowledge the invaluable discussions and support from Dr. Gülçin Pekkurnaz and her laboratory members. We also thank Drs. Susan Ackerman and Byungkook Lim for their insightful comments on the manuscript. We thank Susan Brenner-Morton for ER81 and Nkx6.2 antibodies. We would like to thank the University of California San Diego (UCSD) IGEM Core, UCSD The Human Embryonic Stem Cell Core, the UCSD Nikon Imaging Center, and the UCSD Neurosciences Imaging Center for their expert assistance. This work was supported by The Natural Sciences and Engineering Council of Canada Fellowship to J.G.; and by K01CA190235, the Kavli Foundation, and the Hellman Foundation Fellowship to S.M.G.

## DECLARATION OF INTERESTS

The authors declare no competing interests.

## Supplemental Information

Table S1. RNA-seq_e13.5 sc_CT vs NFIAcKO_DEGs

Table S2. ChIP-seq_e13.5 sc_NFIA_results

Table S3. Table S3 ATAC-seq_CT e13.5 motor neuron promoter_results

Table S4. RNA-ChIP-ATAC overlap-seq_135.e motor neurons_DEGs Table S5. Primer table

## SUPPLEMENTAL FIGURES

**Figure S1:**
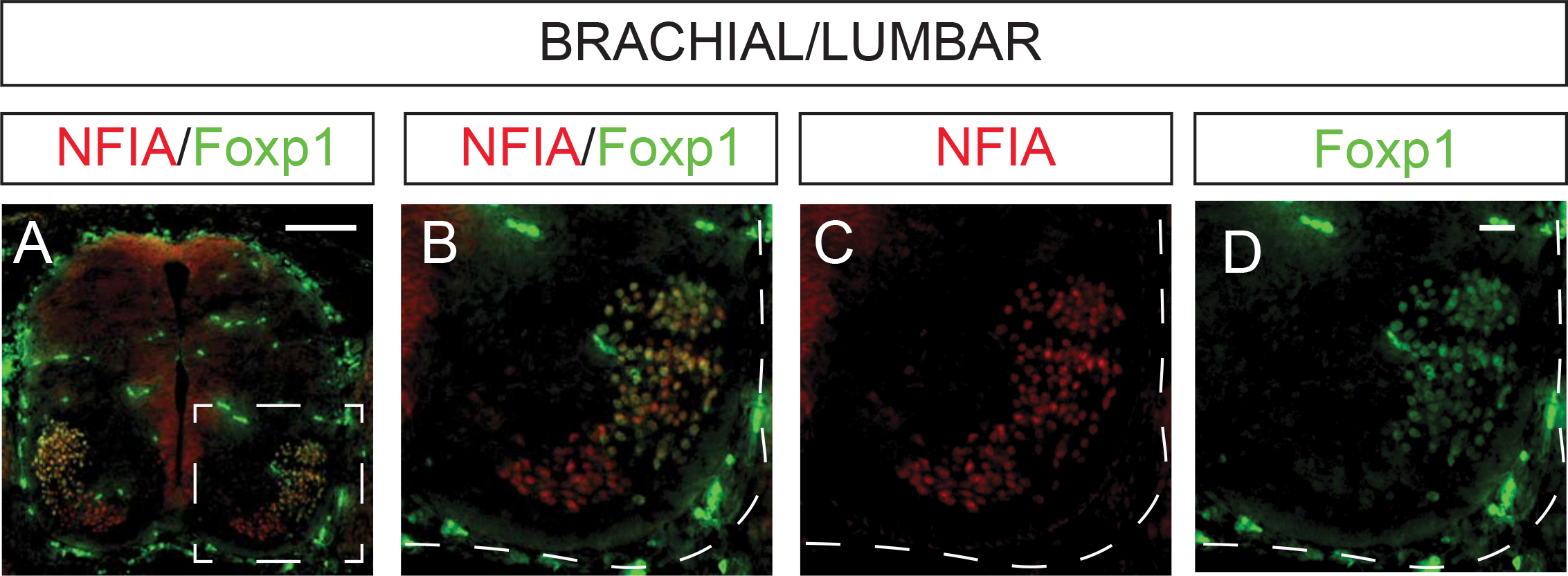
NFIA is expressed in mouse lateral motor columns (LMC) (A) Immunofluorescent staining of NFIA (red) and Foxp1 (green) of E13.5 control mouse spinal cord. (B) Higher magnification image of boxed region in (A). (C) Immunofluorescent staining of NFIA. (D) Immunofluorescent staining of Foxp1. Scale bars represent 100 μm.

**Figure S2:**
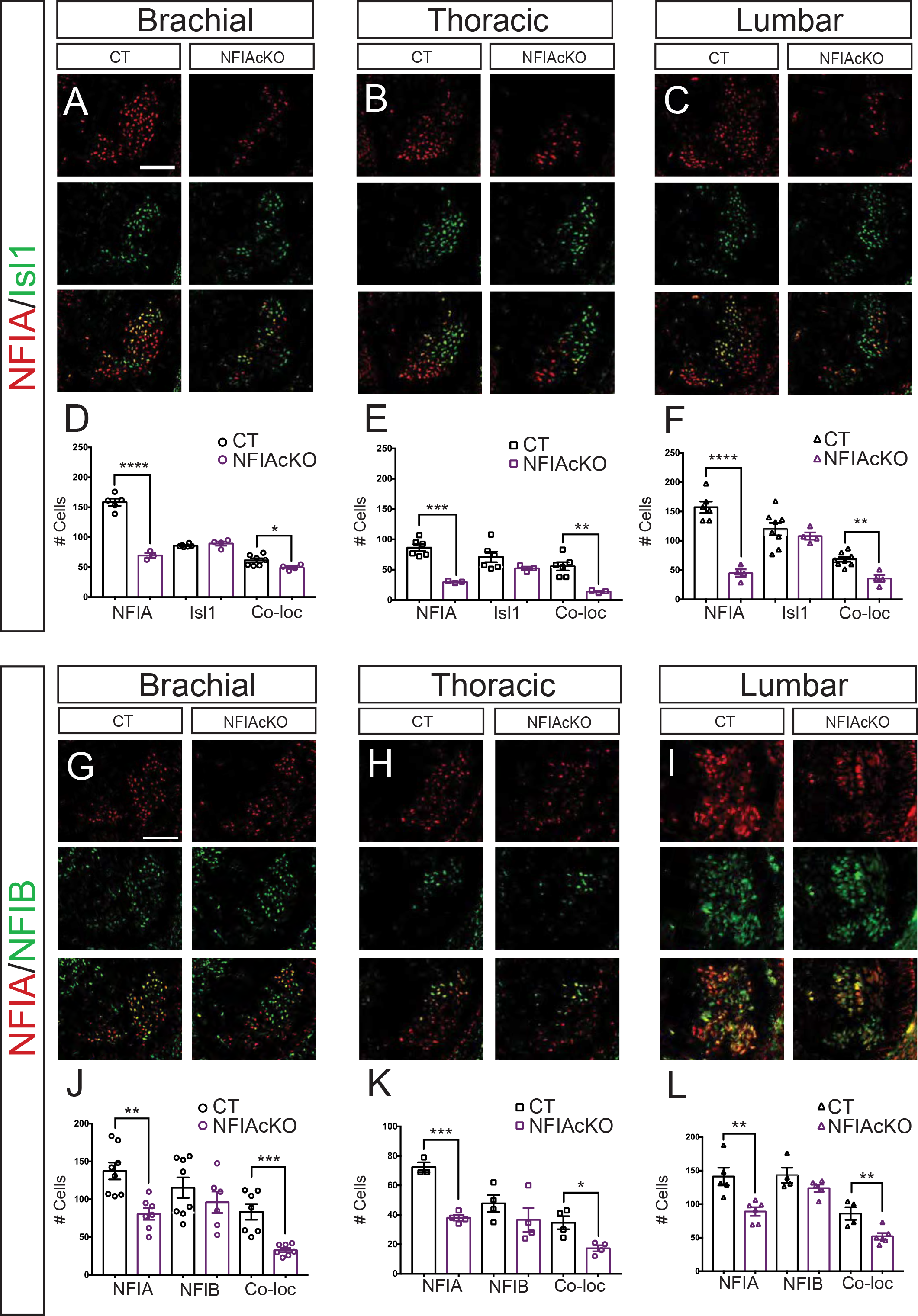
Characterization of NFIA expression in NFIA conditional knockout (NFIAcKO) mice (A-C) Immunofluorescent staining of NFIA (red) and Isl1 (green) on E13.5 control (CT) and NFIAcKO ventral spinal cords at brachial (A), thoracic (B), and lumbar (C) levels. (D-F) Quantification of NFIA-positive, Isl1-positive, and NFIA/Isl1 co-positive (Co-loc) cells showing that NFIA is markedly decreased in NFIAcKO mice at brachial (D), thoracic (E), and lumbar (F) levels. (n=3-8) (G-I) Immunofluorescent staining of NFIA and NFIB on E13.5 CT and NFIAcKO ventral spinal cords at brachial (G), thoracic (H), and lumbar (I) levels. (J-L) Quantification of NFIA-positive, NFIB-positive, and NFIA/NFIB co-loc cells showing no change in NFIB expression at brachial (J), thoracic (K), and lumbar (L). For all images, right ventral quadrant of the spinal cord is shown. Scale bars represent 100 μm. All data are represented as mean ± SEM. p* ≤0.05; p** ≤0.01, p***≤0.001 , p****≤0.0001 by Student’s T- test.

**Figure S3.**
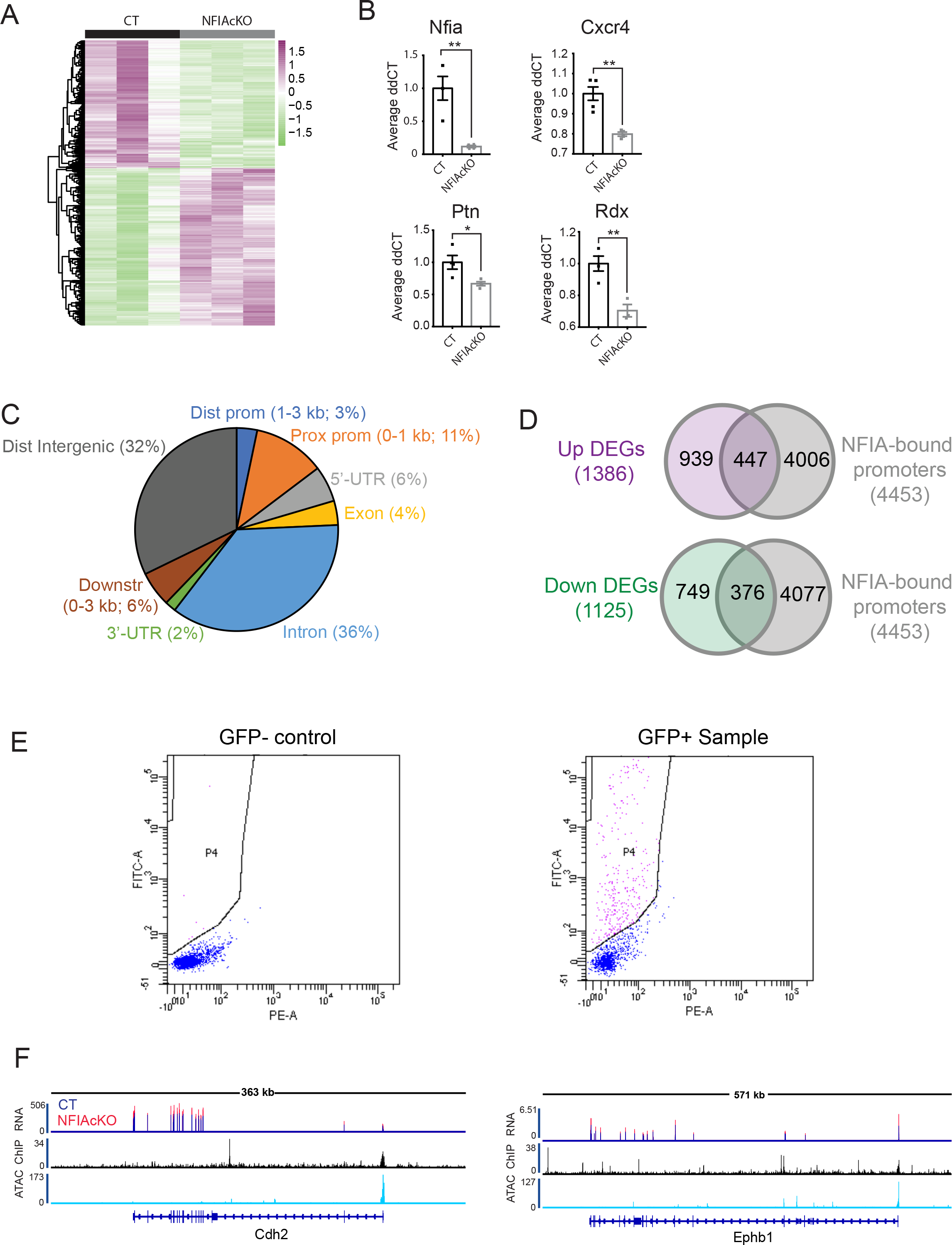
RNA-seq, ChIP-seq, and ATAC-seq analysis of (CT) and NFIA conditional knockout (NFIAcKO) spinal cords (A) Hierarchical clustering of bulk samples based on normalized gene expression in CT and NFIAcKO spinal cords. (B) RT-qPCR analysis confirming decrease of Nfia abundance in NFIAcKO spinal cords and validation of Cxcr4, Ptn, Rdx mRNA in CT and NFIAcKO spinal cords. Values were normalized to Cyclophilin mRNA abundance and expressed as fold change relative to CT (n=3-4). (C) Genomic location of NFIA binding determined by ChIP-seq. (D) Venn diagram of overlap between genes with NFIA-bound promoters (ChIP-seq) and differentially expressed genes (DEGs; RNA-seq) (E) Representative fluorescence activated cell sorting (FACS) plots showing the gating strategy for GFP+ cells isolated from E13.5 spinal cords. Negative control (lacking GFP expression) is on the left. (F) RNA-seq, ATAC-seq, and ChIP-seq gene tracks around the Cdh2 (left) and Ephb1 (right) loci. All data are represented as mean ± SEM. p* ≤0.05; p** ≤0.01 by Student’s T-test.

**Figure S4:**
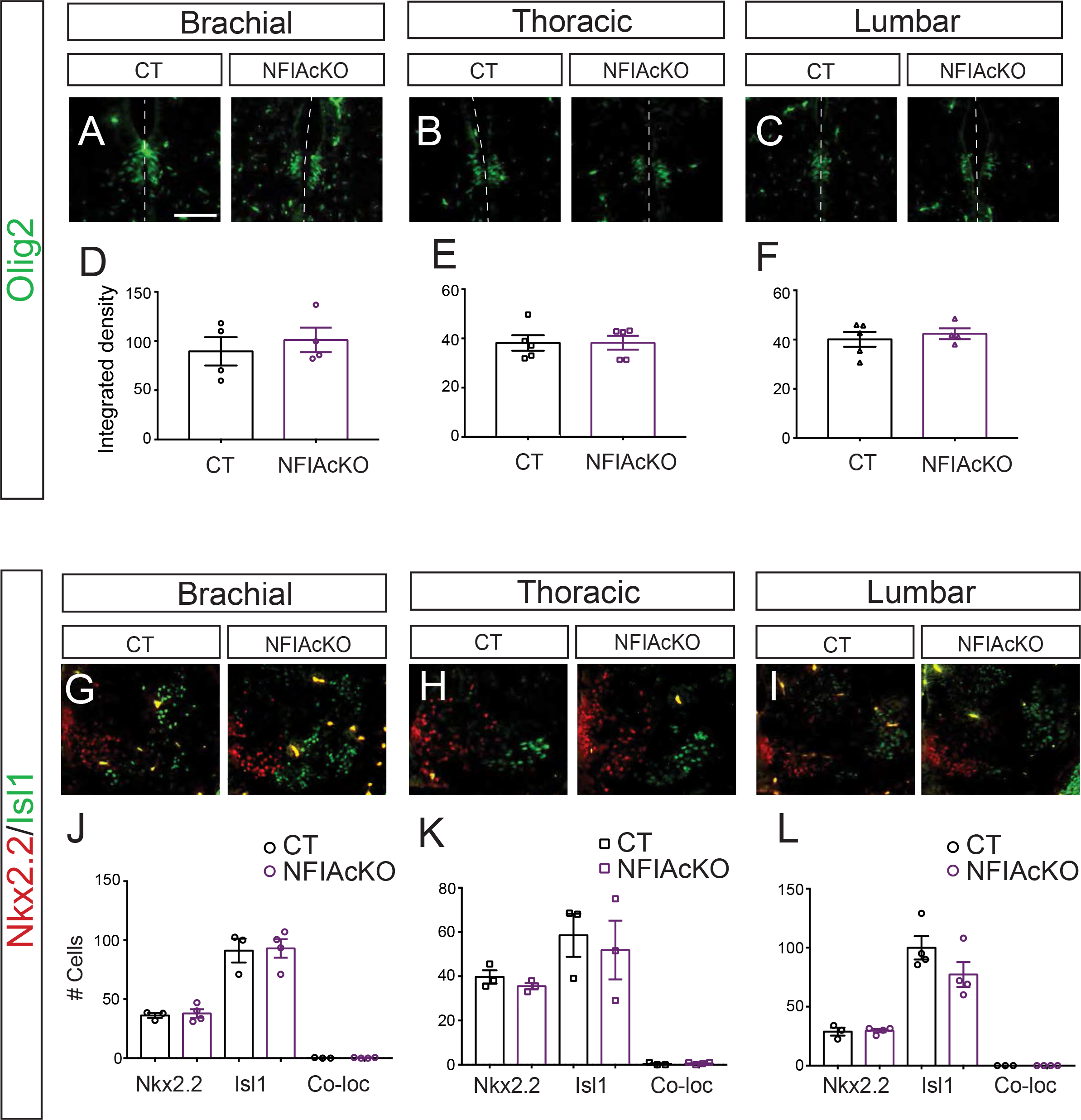
NFIA conditional knockout (NFIAcKO) does not affect ventral spinal cord progenitor cells (A-C) Immunofluorescent staining of Olig2 on E13.5 control (CT) and NFIAcKO ventral spinal cords at brachial (A), thoracic (B), and lumbar (C) levels. (D-F) Quantification of Olig2 in brachial (D), thoracic (E), and lumbar (F) regions. (n=4-5) (G-I) Immunofluorescent staining of Nkx2.2 (red) and Isl1 (green) on E13.5 CT and NFIAcKO ventral spinal cords at brachial (G), thoracic (H), and lumbar (I) levels. (J-L) Quantification of Nkx2.2-positive, Isl1-positive, and Nkx2.2/Isl1 co-positive (Co-loc) cells. (n=3-4). Scale bars represent 100 μm. All data are represented as mean ± SEM.

**Figure S5:**
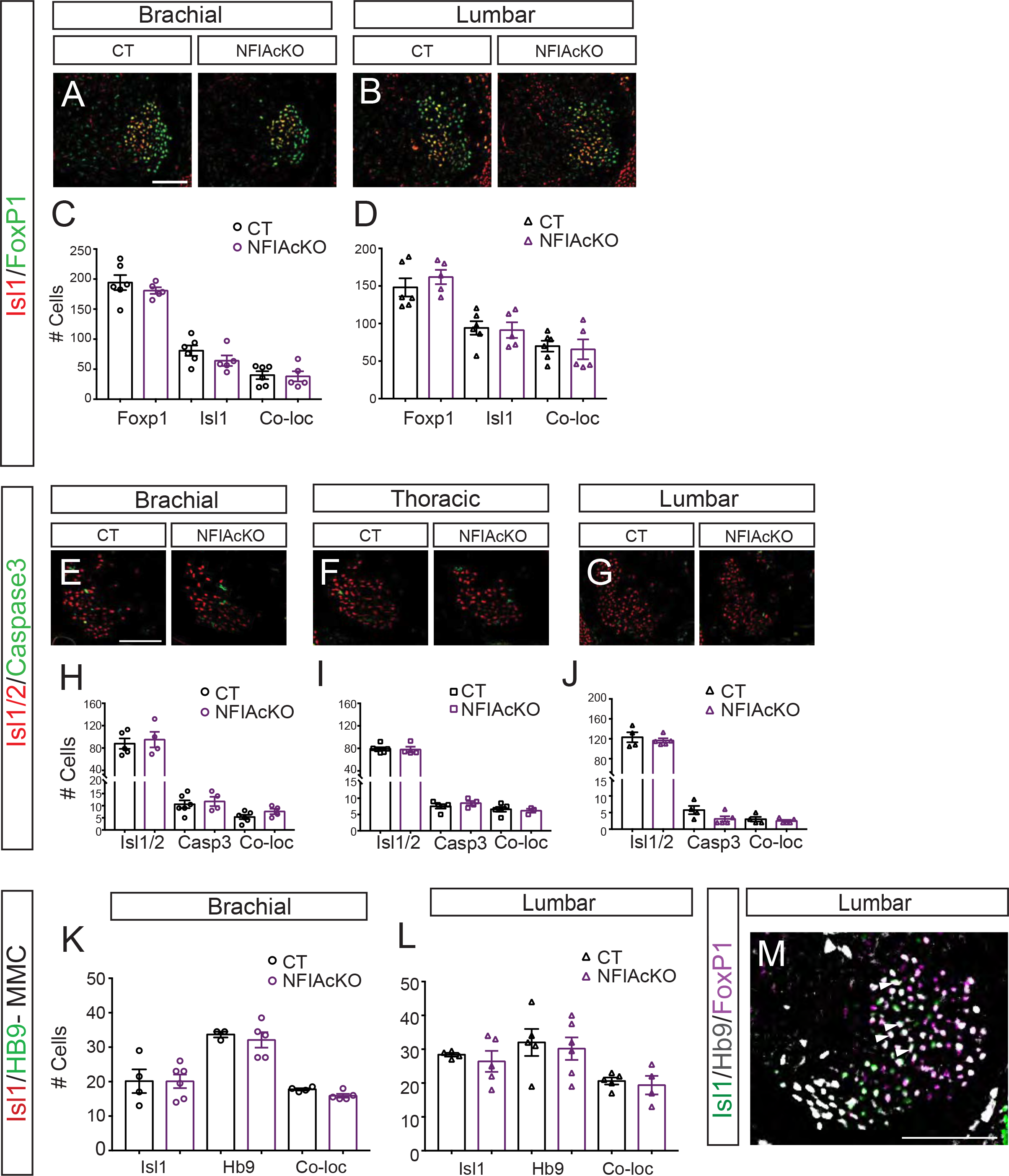
Total lateral motor culmns (LMC) motor neuron numbers are unchanged in NFIA conditional knockout (NFIAcKO) mice (A-B) Immunofluorescent staining of LMC markers Isl1 (red) and Foxp1 (green) on E13.5 control (CT) and NFIAcKO ventral spinal cords at brachial (A) and lumbar (B) levels. (C-D) Quantification of Isl1-positive, Foxp1-positive, and Isl1/Foxp1 co-positive (Co-loc) cells in brachial (C) and lumbar (D) regions. (n=3-6) (E-G) Immunofluorescent staining of activated caspase 3 (cell death marker; green) and Isl1/2 (motor neuron marker; red) on E13.5 CT and NFIAcKO ventral spinal cords at brachial (E), thoracic (F), and lumbar (G) levels. (H-J) Quantification of Isl1/2-positive, Caspase3-positive, and Isl1/2 & Caspase3 Co-loc cells in brachial (H), thoracic (I), and lumbar (J) regions. (n=4-6) (K-L) Quantification of median motor column (MMC) markers Isl1 and Hb9 in E13.5 CT and NFIAcKO spinal cords at brachial (K) and lumbar (L) spinal cord levels. For all images, right ventral quadrant of the spinal cord is shown. Scale bars represent 100 μm. All data are represented as mean ± SEM. (M) Immunofluorescent staining of Isl1 (green), Hb9 (grey) and Foxp1 (magenta) on E13.5 NFIAcKO ventral spinal cords at lumbar level. Arrowheads denote Isl1/Hb9/Foxp1 triple-positive cells in the LMC.

**Figure S6:**
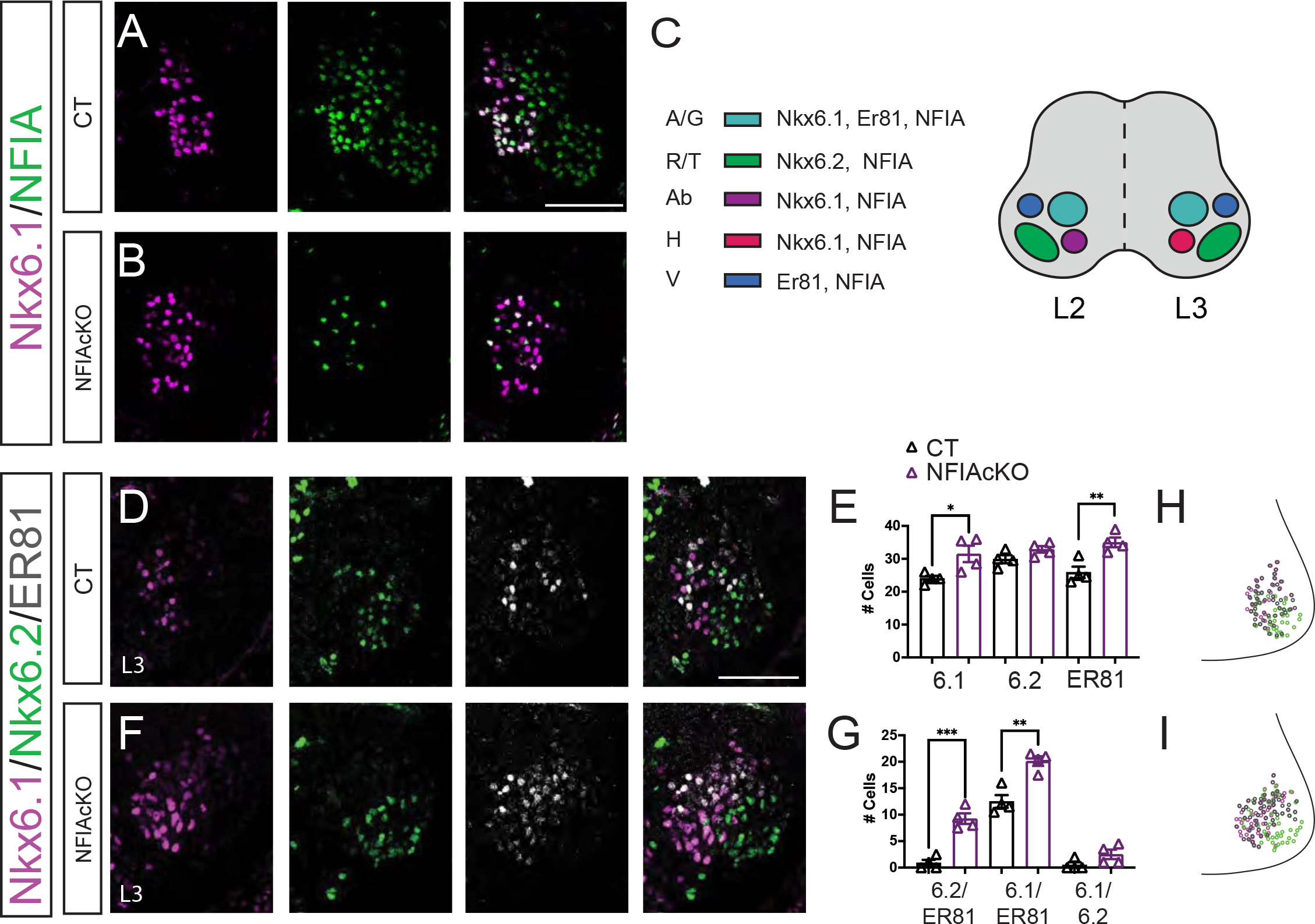
Lateral motor colmn (LMC) segregation defects of L3 motor pools in NFIA mutants (A-B) Immunofluorescent staining of NFIA (green) and Nkx6.1 (magenta) at L2 regions of the E13.5 ventral spinal cord in control (CT) and NFIA conditional knockout (NFIAcKO) embryos. (C) Schematic of L2 and L3 mouse motor pools. At L2 Nkx6.1+/Er81+ demarcates the adductor/gracilis (A/G)-innervating pools, Nkx6.2 marks the femoris/tensor fasciae latae (R/T)-innervating pool, the adductor brevis (Ab)-innervating pool can be defined by Nkx6.1, and the vasti (V)-innervating pool is labeled by Er81 expression. At L3 the Ab-innervating is no longer demarcated by Nkx6.1. Instead, Nkx6.1 defines the hamstring (H)-innervating pool. (D,F) Immunofluorescent staining of motor pool markers Nxk6.1/Nkx6.2/Er81 at L3 lumbar level in E13.5 CT (D) and NFIAcKO (F) spinal cords. (E,G) Quantification of L3 neurons in CT and NFIAcKO lumbar spinal cords. (n=4-5) (H-I) Digitally reconstructed distribution of CT (H) and NFIAcKO (I) lumbar L3 motor pools, shown as a transverse projection (dot plot). For all images, right ventral quadrant of the spinal cord is shown. Scale bars represent 100 μm. All data are represented as mean ± SEM. p* ≤0.05; p** ≤0.01, p***≤0.001 by Student’s T-test.

**Figure S7:**
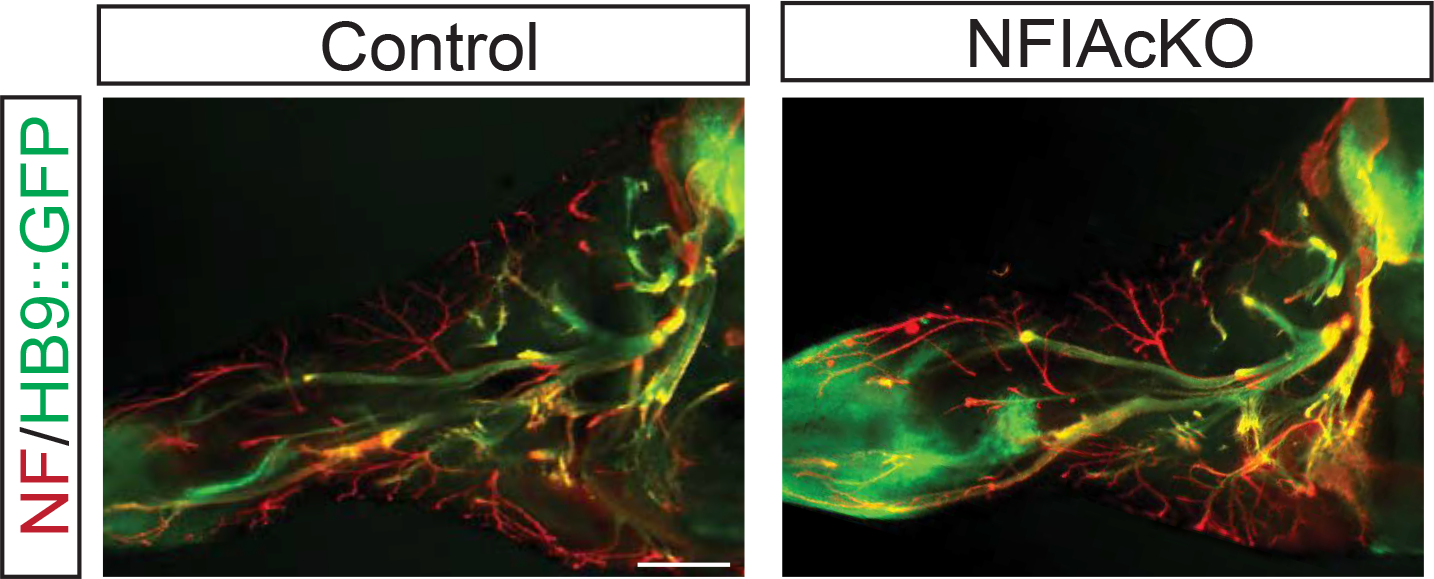
Lateral motor column (LMC) projections from the base of the limb are unaffected by loss of NFIA. Flat-mount neurofilament (NF; red) and HB9::GFP staining of hindlimb innervation patterns in control NFIA^ff^;Hb9::GFP mice and NFIA conditional knockout (NFIA^ff^;Nestin-Cre;Hb9::GFP; NFIAcKO) at E13.5.

**Figure S8:**
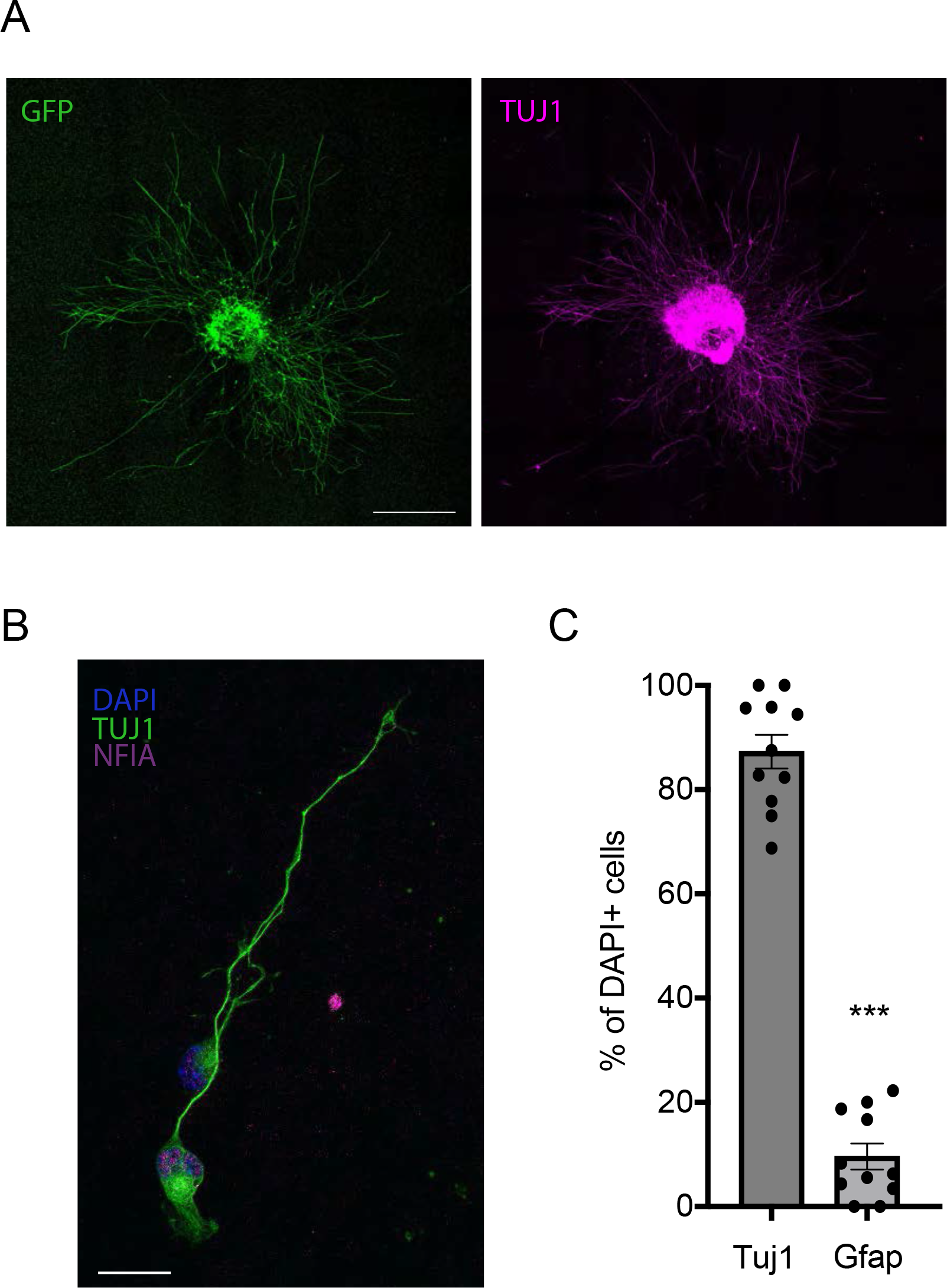
Primary motor neuron cultures contain high numbers of neurons and nominal astrocytes (A) Immunofluorescent staining of GFP (green) and Tuj1 (magenta) in control explant cultures from E13.5 brachial spinal cord after 4 DIV. Scale bar represents 500 μm. (B) Immunofluorescent staining of NFIA (magenta), Tuj1 (green) and DAPI (blue) in 3 DIV mouse primary motor neuron cultures. (B) Quantification of Tuj1+ and Gfap+ cells in control primary motor neuron cultures. (n=11) Scale bar represents 20 μm. All data are represented as mean ± SEM. p***≤0.001 by Student’s T-test.

## Notes

### Competing Interest Statement

The authors have declared no competing interest.

